# Mechanical opening of the αE-catenin M-region gates afadin condensation in cardiomyocytes

**DOI:** 10.64898/2026.09.27.754825

**Authors:** Sabine Pokutta, Yerin Han, Korina I. Karpov, Jonathon A. Heier, William I. Weis, Adam V. Kwiatkowski

## Abstract

In mechanically active tissues, cell-cell adhesions must withstand high and dynamic loads to maintain tissue integrity. The ability to detect and withstand load centers on the mechanosensitive adherens junction (AJ), which couples the actin networks of adjacent cells. The mechanosensor αE-catenin responds to force by opening its Middle (M) region, revealing cryptic binding sites for adaptor proteins such as afadin and vinculin. Here we show that the afadin coiled-coil (CC) region binds the open αE-catenin M-region with modest affinity and fast exchange, unlike the high-affinity, long-lived binding of vinculin, suggesting distinct roles. In cardiomyocytes, the afadin CC is necessary and sufficient for afadin localization at high-load AJs, where it exhibits the dynamic, hexanediol-sensitive properties of a biomolecular condensate. By contrast, the afadin CC is dispensable for recruitment at low-load epithelial AJs, suggesting that junctional load determines the basis of afadin recruitment. We propose that force-gated opening of αE-catenin seeds afadin condensate formation, a mechanism that drives reorganization of high-load AJs.

## Introduction

The adherens junction (AJ) physically links the actin networks of adjacent cells, and both bears and responds to the load transmitted across them (Charras and Yap, 2018; Leckband and de Rooij, 2014; Lenne et al., 2021; Mege and Ishiyama, 2017; Noordstra et al., 2023). The cadherin-catenin complex forms the AJ core (Harris and Tepass, 2010; Meng and Takeichi, 2009). Classical cadherins are transmembrane proteins that bind homotypically to cadherins on opposing membranes to link cells (Pokutta and Weis, 2007; Shapiro and Weis, 2009). Cadherin-mediated adhesion requires cytoplasmic linker proteins called catenins that couple cadherins to the actin cytoskeleton (Buckley et al., 2014; le Duc et al., 2010; Yao et al., 2014; Yonemura et al., 2010). Yet how mechanical load controls adaptor recruitment, and how this recruitment reinforces adhesions under load, remains unclear.

αE-catenin transduces load through two distinct mechanisms: the actin-binding domain (ABD), which forms a catch bond with F-actin (Arbore et al., 2022; Buckley et al., 2014; Xu et al., 2020), and force-dependent opening of the mechanosensitive Middle (M) region (Choi et al., 2012; Pokutta et al., 2002; Yao et al., 2014). M-region opening relieves autoinhibition to reveal cryptic binding sites for adaptor proteins such as vinculin and afadin. Vinculin, a mechanosensitive actin-binding protein similar to αE-catenin, is recruited to the force-opened M-region, creating an additional link to actin filaments (Huang et al., 2017; le Duc et al., 2010; Thomas et al., 2013; Yonemura et al., 2010). Afadin is a multidomain scaffold protein that is recruited to junctions via binding to force-activated αE-catenin as well as interactions with nectin transmembrane proteins (Mandai et al., 1997; Pokutta et al., 2002; Sakakibara et al., 2020; Tachibana et al., 2000; Takai and Nakanishi, 2003). Afadin also assembles into biomolecular condensates that concentrate junctional components and reinforce adhesion (Kuno et al., 2025). Because condensate formation is inherently concentration-dependent, force-dependent recruitment could control both afadin localization and condensation at junctions. Whether mechanical load gates afadin condensate formation at AJs, however, remains unknown.

The molecular basis of afadin function at AJs has become clearer with recent biochemical and structural advances. The αE-catenin binding site in afadin was mapped to the N-terminal stretch of the afadin coiled-coil (CC) region (Maruo et al., 2018), which is embedded within a large C-terminal intrinsically disordered region (IDR) (Gurley et al., 2023; Kuno et al., 2025). Afadin CC binding to the αE-catenin/β-catenin complex *in vitro* increases the affinity of αE-catenin for F-actin (Sakakibara et al., 2020), suggesting a role for afadin in regulating the AJ-actin interface. A subsequent cryo-EM study revealed that the C-terminal region of the CC binds and bridges adjacent αE-catenin ABDs along actin filaments to promote cooperative binding (Gong et al., 2025). Thus, the afadin CC contains two separate binding sites for αE-catenin. Afadin also forms condensates at epithelial AJs, driven by multivalent interactions and the afadin IDR (Kuno et al., 2025). Similarly, the IDR of the fly ortholog Canoe mediates junctional localization (Jensen et al., 2025). Thus, afadin acts at epithelial junctions through at least two domains: the CC region that binds αE-catenin to reinforce the AJ–actin interface, and the IDR that drives condensate formation and locally concentrates afadin.

In cardiac muscle, the AJ anchors contractile myofibrils at highly specialized cardiomyocyte junctions called intercalated discs (Vite and Radice, 2014). To build and maintain AJs under high load, cardiomyocytes recruit a large set of actin-binding and adaptor proteins that strengthen the connection between the cadherin-catenin complex and F-actin, including vinculin and afadin (Li et al., 2019). Our previous work showed that force-activated αE-catenin is required to recruit both vinculin and afadin to cardiomyocyte AJs, and that vinculin binding to αE-catenin was necessary and sufficient for myofibril coupling (Merkel et al., 2019). This is consistent with recent *in vitro* studies that showed that, within the E-cadherin/β-catenin/αE-catenin/vinculin complex, vinculin is the primary connection to actin filaments (Gong et al., 2025). In contrast, afadin recruitment to αE-catenin in the absence of vinculin in cardiomyocytes promoted cell-cell adhesion but failed to support myofibril coupling at AJs (Merkel et al., 2019). These data suggested that vinculin provides needed stability to the AJ under load, whereas afadin serves a distinct function.

Building on our past work(Merkel et al., 2019), here we define the molecular basis of afadin recruitment and show that junctional afadin displays the dynamic, hexanediol-sensitive properties of a biomolecular condensate. When binding to the open αE-catenin M-region, afadin CC exhibits modest affinity and rapid kinetics, whereas vinculin forms a high-affinity, long-lived complex, suggesting distinct functional roles. Afadin CC and vinculin can bind the M-region simultaneously and relieve αE-catenin autoinhibition *in vitro*. The second site in the afadin CC that engages the actin-bound αE-catenin ABD does not detectably increase afadin’s affinity for αE-catenin, indicating that M-region opening is the principal determinant of afadin recruitment to αE-catenin. In cardiomyocytes, the afadin CC is necessary and sufficient for junctional recruitment, and actomyosin contractility is required to maintain afadin localization. In contrast, afadin CC is not recruited to low-load junctions in epithelial MDCK cells. We propose a force-gated condensate mechanism at high-load AJs, in which mechanical opening of the αE-catenin M-region seeds afadin condensate formation to regulate AJ organization and reinforcement.

## Results

### Autoinhibition regulates αE-catenin binding to afadin

To define the molecular basis and regulation of the afadin–αE-catenin interaction, we first performed *in vitro* biochemistry using recombinant afadin and αE-catenin constructs (Fig. 1). αE-catenin contains an N-terminal β-catenin-binding domain, a Middle (M) region containing three domains (M1, M2, M3) that together provide binding sites for adaptor proteins, and a C-terminal F-actin-binding domain (ABD) (Fig. 1, A). Afadin is a large multidomain protein with N-terminal RA1-RA2, FHA, DIL, and PDZ domains followed by a C-terminal coiled-coil region embedded in an extensive intrinsically disordered region (IDR) (Fig. 1, B).

**Figure 1.**
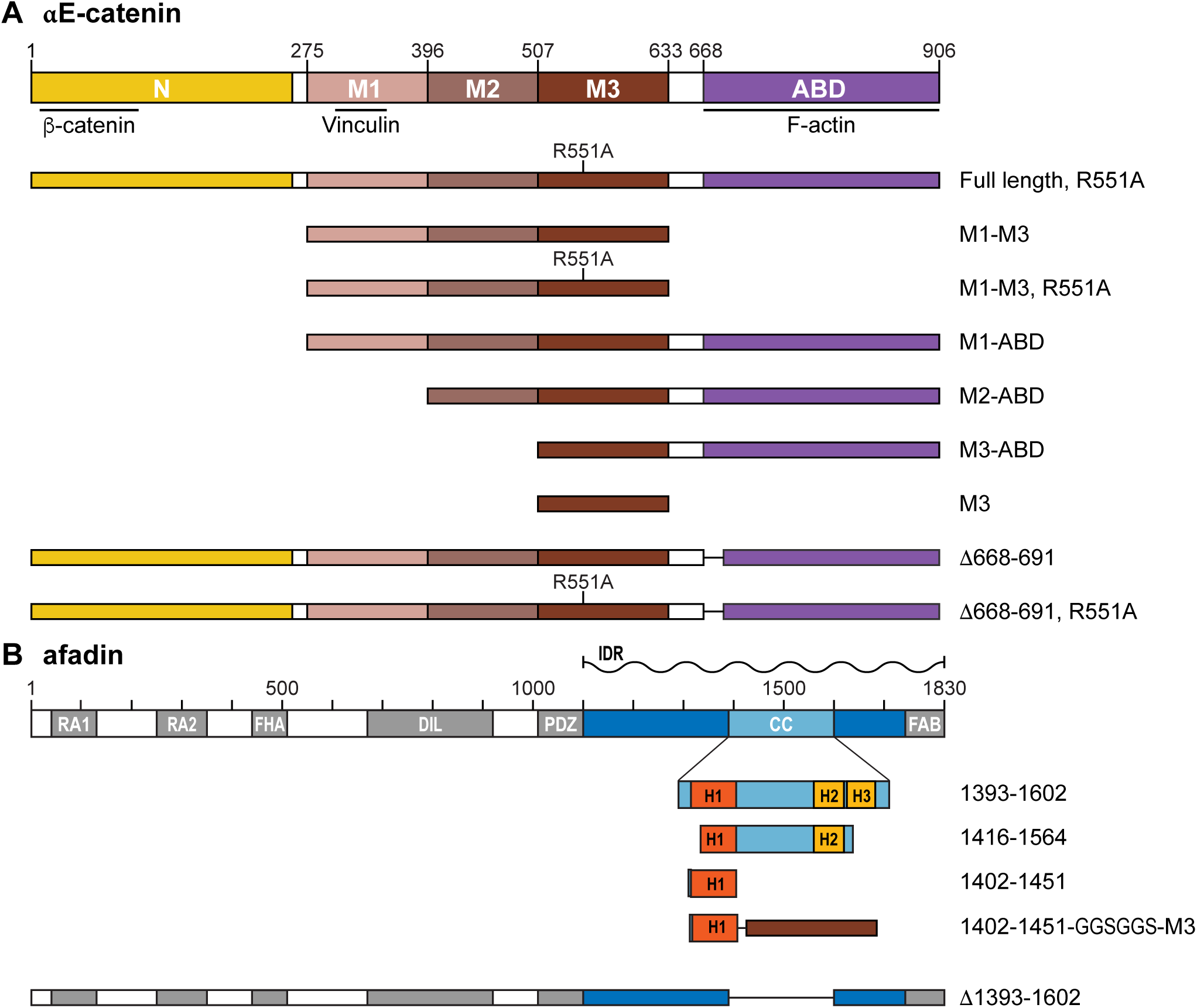
αE-catenin and afadin domain organization and constructs used in this study. **(A)** Cartoon schematic of αE-catenin domain organization. N-terminal region (N), Middle(M)-region domains (M1, M2, M3), and actin-binding domain (ABD) are labeled. Binding sites for β-catenin, vinculin, and F-actin are indicated. αE-catenin constructs used in ITC experiments, actin cosedimentation assays, and cell biology experiments are listed below. R551A point mutation relieves M-region autoinhibition. **(B)** Cartoon schematic of afadin domain organization. Ras-associating (RA1 and RA2), forkhead-associated (FHA), Dilute (DIL), PDZ, coiled-coil (CC), and F-actin binding (FAB) domains are labeled. C-terminal intrinsically disordered region (IDR) is marked. Constructs used in *in vitro* and cell biology assays are listed below. The CC region contains three alpha helices (H1, H2, H3).

Previous studies have shown that αE-catenin binding to afadin is regulated by autoinhibition (Sakakibara et al., 2020). The afadin binding site was mapped by pull-down assays to the M3 domain of αE-catenin and to residues 1400-1460 of afadin (Maruo et al., 2018; Sakakibara et al., 2020). To quantitatively characterize how αE-catenin autoinhibition regulates afadin binding, we performed ITC. We used two afadin constructs for our binding experiments: the afadin CC region (aa 1393-1602), based on a mouse afadin fragment previously shown to bind αE-catenin (Sakakibara et al., 2020), and a smaller fragment, afadin 1416-1564, which showed better expression and which we used in our initial mapping studies. Comparable affinities were observed with both afadin constructs (1416-1564 and 1393-1602), indicating that the core binding region is contained within residues 1416-1564 (Fig. 2; Fig. S1).

**Figure 2.**
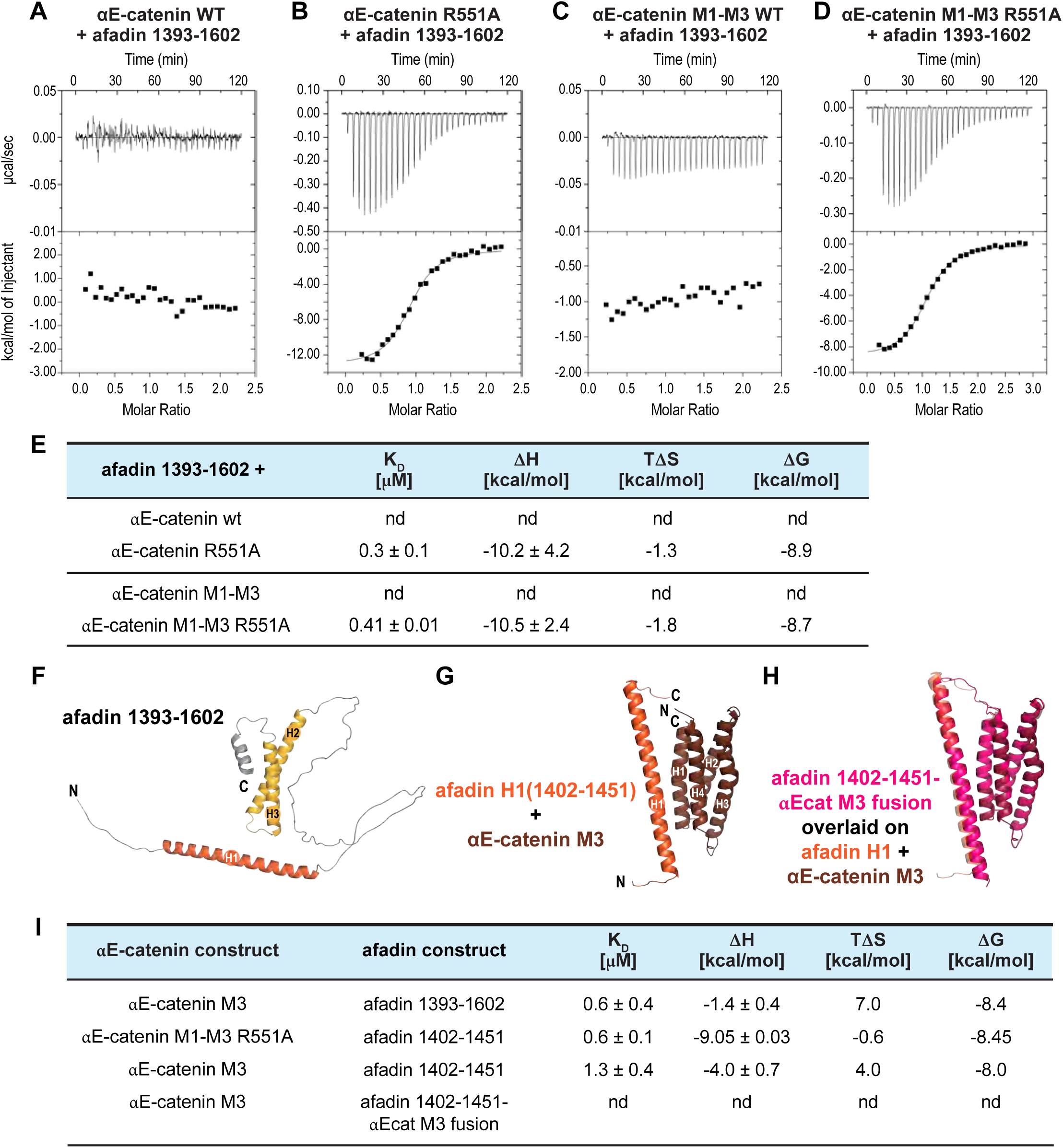
αE-catenin M-region autoinhibition prevents afadin binding. **(A-D)** Representative ITC traces for afadin 1393-1602 binding to wild-type or R551A αE-catenin full-length (A, B) or M1-M3 protein (C, D). **(E)** Thermodynamic parameters derived from duplicate measurements; nd = not detected. **(F)** AlphaFold predicted structure of afadin residues 1393-1602 with N- and C-terminal helical regions colored in orange (H1) and yellow (H2, H3). **(G)** AlphaFold-Multimer predicted structure of the complex formed between afadin 1400-1458 and αE-catenin 507-635 (M3 domain). Afadin H1 is colored orange; αE-catenin M3 domain helices H1-H4 are colored brown. **(H)** Superposition of the predicted structures of the afadin/αE-catenin M3 complex (same color scheme as in (G)) and the afadin 1402-1451-GGSGGS-αE-catenin 507-633 fusion protein. The fusion protein is shown in pink. **(I)** Thermodynamic parameters derived from ITC measurements of binding between αE-catenin M3 and afadin 1393-1602, αE-catenin M1-M3 R551A or αE-catenin M3 and afadin H1 (1402-1451) and αE-catenin M3 and afadin 1402-1451-αE-catenin M3 fusion protein. When binding was observed, thermodynamic parameters were derived from duplicate measurements; nd = not detected.

No binding was detected between afadin and full-length αE-catenin (Fig. 2, A and E) or αE-catenin/β-catenin 78-671 complex (Fig. S1, A and G), or between afadin and the αE-catenin M1-M3 fragment (Fig. 2, C and E). The tertiary structure of the αE-catenin M-region is stabilized by a salt-bridge network, which can be disrupted by mutating Arg 551 to Ala (R551A) (Ishiyama et al., 2013). As shown by ITC, the β-catenin 78-671/αE-catenin R551A complex (Fig. S1, B and G), full-length αE-catenin R551A (Fig. S1, C and G; Fig. 2, B and E), and the αE-catenin M1-M3 R551A construct (Fig. 2, D and E) all bound afadin with 0.3–0.6 μM affinity, demonstrating that the R551A mutation relieves autoinhibition for afadin binding.

These results confirm that αE-catenin binding to afadin is regulated by autoinhibition, and that destabilization of the M-region salt-bridge network releases autoinhibition. Notably, we show that afadin binds with modest affinity to the αE-catenin M-region. This contrasts with vinculin, which binds with low affinity (2 μM, (Choi et al., 2012)) to the autoinhibited M-region and with high affinity to the open M-region (4 nM, (Terekhova et al., 2019)).

### The αE-catenin M3 domain interacts with the H1 helix of the afadin coiled-coil region

To define the minimal binding interface between afadin and the αE-catenin M-region, we performed ITC with a series of αE-catenin deletion constructs. Similar to the R551A mutation, deletion of the M1 domain relieves autoinhibition (Sakakibara et al., 2020), and the αE-catenin M2-ABD fragment, M3-ABD and the isolated M3 domain bound afadin 1416-1564 (Fig. S1, D-G) or afadin 1393-1602 (Fig. S1, H; Fig. 2, I) with affinities ranging between 0.6 and 1.0 μM. Thus, the M2 and ABD domains do not contribute substantially to binding beyond the M3 domain alone, and the binding site for afadin is therefore predominantly contained within the M3 domain.

The AlphaFold3 model of afadin 1393-1602 predicts two helical regions within the αE-catenin-binding region: a single extended helix (H1) located at the N-terminus (1406-1446) and two helices (H2 and H3) connected through a short loop at the C-terminus (1520-1581) (Fig. 2, F). We found that the affinity of the N-terminal helical region (afadin 1402-1451) for αE-catenin M1-M3 R551A was 0.6 μM (Fig. S1, I; Fig. 2, I), comparable to that of afadin 1393-1602 (Fig. 2, D and E). Using the isolated M3 domain of αE-catenin and afadin 1402-1451 resulted in only slightly weaker binding (K_D_ = 1.3 μM) (Fig. S1, J; Fig. 2, I), demonstrating that these fragments recapitulate most of the binding interface. In the AlphaFold-Multimer predicted structure of the M3 domain/afadin 1402-1451 complex, H1 of afadin packs against helices 1 and 4 of the M3 domain four-helix bundle (Fig. 2, G). Lys 1417 of the afadin H1 helix is predicted to form the first N-terminal contact with the M3 domain—consistent with the observation that the shorter afadin fragment (1416-1564) and the longer afadin 1393-1602 bind with comparable affinities (Fig. 2; Fig. S1).

Taking the predicted orientation of the afadin H1 helix into account, we generated an N-terminal afadin 1402-1451–αE-catenin M3 fusion protein in which both fragments are connected through a six-amino acid linker. The predicted structure of the fusion protein superimposes well with the predicted complex (Fig. 2, H). This construct did not interact with free αE-catenin M3 by ITC (Fig. S1, K; Fig. 2, I), suggesting that the afadin 1402-1451 fragment is interacting intramolecularly with the M3 domain to which it is fused and is therefore unavailable for binding to free M3. This result is consistent with the AlphaFold-Multimer prediction and supports the predicted binding orientation.

### Afadin and vinculin can simultaneously bind αE-catenin and mutually relieve autoinhibition

Vinculin and afadin bind to distinct domains within the M-region: vinculin binds to the M1 domain (Choi et al., 2012; Rangarajan and Izard, 2012), whereas afadin binds primarily to M3 ((Sakakibara et al., 2020), our data). As shown previously in pull down assays, vinculin and afadin can bind simultaneously to αE-catenin M319G/R326E/R551E (Gong et al., 2025), and when combined in equimolar concentrations, afadin 1393-1602, vinculin 1-839 (in which deletion of the C-terminal ABD relieves autoinhibition), and αE-catenin M1-M3 R551A form a ternary complex that eluted as a single peak off a SEC column (Fig. 3, A). Afadin bound to the αE-catenin M1-M3 R551A/vinculin complex with 0.5 μM affinity as shown by ITC (Fig. 3, C and F), demonstrating that afadin binding to αE-catenin is not affected by vinculin.

**Figure 3.**
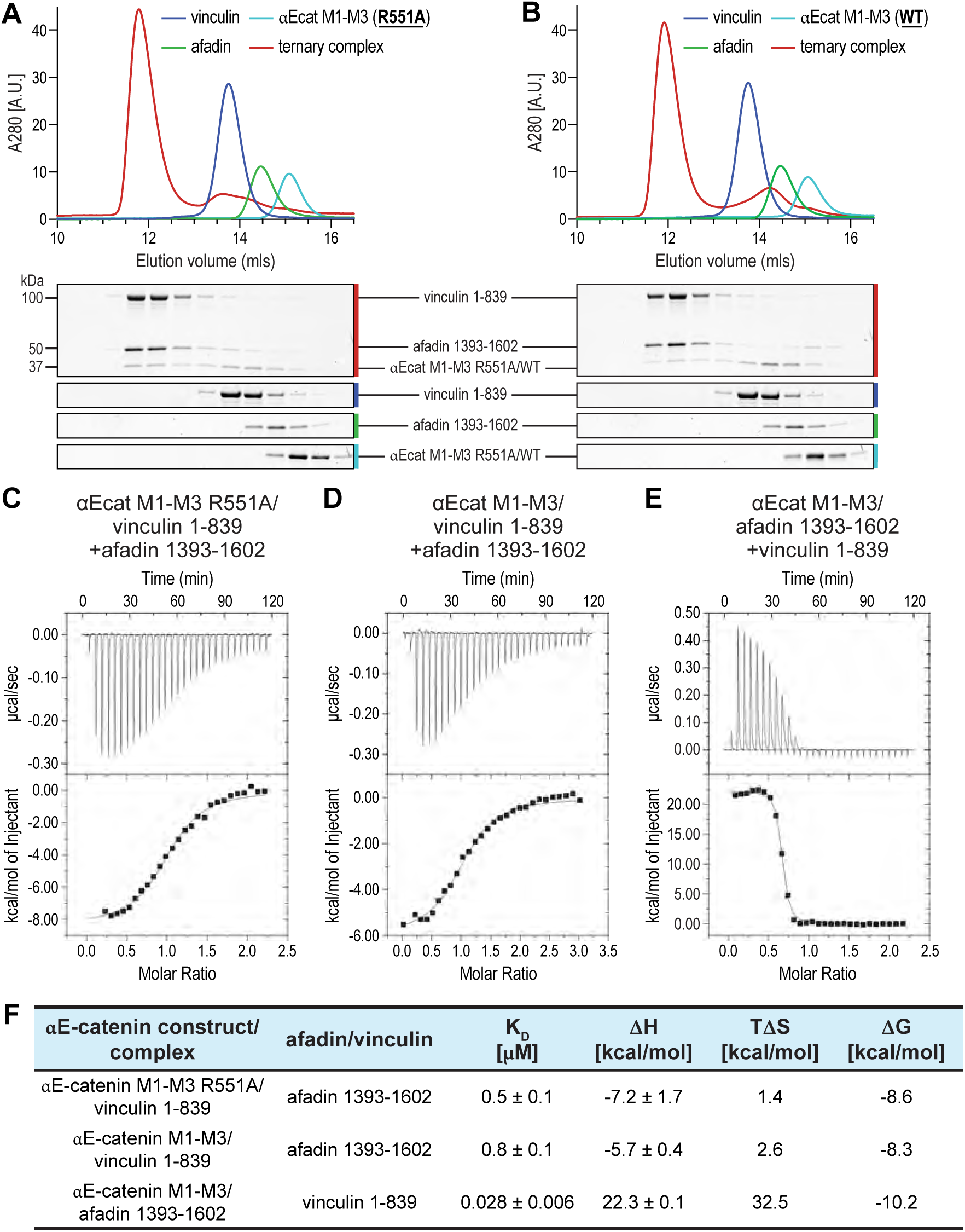
Afadin and vinculin form a ternary complex with αE-catenin by mutually relieving M-region autoinhibition. **(A, B)** Size exclusion chromatograms of equimolar mixtures (ternary complex, red line) of afadin 1393-1602 and vinculin 1-839 with either αE-catenin M1-M3 R551A (A) or αE-catenin M1-M3 (B). For comparison, elution profiles for each individual protein are shown. Afadin 1393-1602 (green line), vinculin 1-839 (dark blue line), αE-catenin (light blue line). Fractions collected from SEC runs were analyzed by SDS-PAGE and Coomassie staining. **(C, D)** Representative ITC traces of afadin 1393-1602 binding to the preformed αE-catenin M1-M3 R551A/vinculin 1-839 complex (C) or αE-catenin M1-M3 in the presence of vinculin 1-839 (D). **(E)** Representative ITC trace of vinculin 1-839 binding to αE-catenin M1-M3 in the presence of afadin 1393-1602. **(F)** Thermodynamic parameters were derived from duplicate measurements.

We further tested whether afadin and vinculin can mutually relieve αE-catenin autoinhibition. αE-catenin M1-M3, which did not interact with afadin in its autoinhibited state (Fig. 2, C and E), nonetheless formed a ternary complex with afadin 1393-1602 and vinculin 1-839 that co-eluted off a SEC column (Fig. 3, B). In the presence of vinculin 1-839, αE-catenin M1-M3 bound afadin 1393-1602 with 0.8 μM affinity (Fig. 3, D and F), whereas no binding was observed with αE-catenin M1-M3 alone. Conversely, in the presence of afadin 1393-1602, vinculin bound to αE-catenin M1-M3 with 28 nM affinity—an approximately 70-fold increase in affinity compared to autoinhibited αE-catenin M1-M3 (Fig. 3, E and F) (Choi et al., 2012). These results demonstrate that afadin and vinculin can mutually relieve αE-catenin autoinhibition.

### Afadin binds actin-associated αE-catenin through two sites with distinct affinities

The recently solved cryo-EM structure of the E-cadherin–β-catenin–αE-catenin–vinculin–afadin CC complex bound to F-actin identified a second αE-catenin binding site within the afadin CC (Gong et al., 2025). This site encompasses the C-terminal afadin helices H2 and H3 that contact both the αE-catenin ABD and F-actin. Afadin binding stabilizes the four-helix bundle conformation of the actin-bound ABD (Gong et al., 2025); which represents the strong binding state of the αE-catenin two state catch bond (Wang et al., 2022; Xu et al., 2020). If and how this additional binding site affects the binding of afadin CC to actin-bound αE-catenin is unknown.

To determine the affinity of afadin for actin-bound αE-catenin, we performed actin cosedimentation assays. To increase the actin-binding affinity of αE-catenin, we introduced a deletion in the ABD (residues 668-691) (Xu et al., 2020) that promotes stable F-actin association while retaining all ABD residues that contact afadin helices H2 and H3 (Gong et al., 2025). This construct bound actin with an affinity of approximately 0.3 μM (Fig. S2). Binding assays were performed with the β-catenin/αE-catenin complex using a β-catenin fragment comprising the αE-catenin binding site (residues 78-151). We measured a dissociation constant of 9.5 μM for afadin binding to actin-bound αE-catenin Δ668-691, in which the M-region is autoinhibited, and binding occurs solely through the actin-bound ABD (site 2) (Fig. 4, A). In contrast, actin-associated αE-catenin Δ668-691 R551A—in which both afadin H1/αE-catenin M3 binding site (site 1) and the binding site in the ABD (site 2) are available—bound afadin with approximately 25-fold higher affinity (K_D_ = 0.38 μM) (Fig. 4, B), consistent with the affinity determined by ITC for the αE-catenin M1-M3 R551A/afadin interaction (Fig. 2, E).

**Figure 4.**
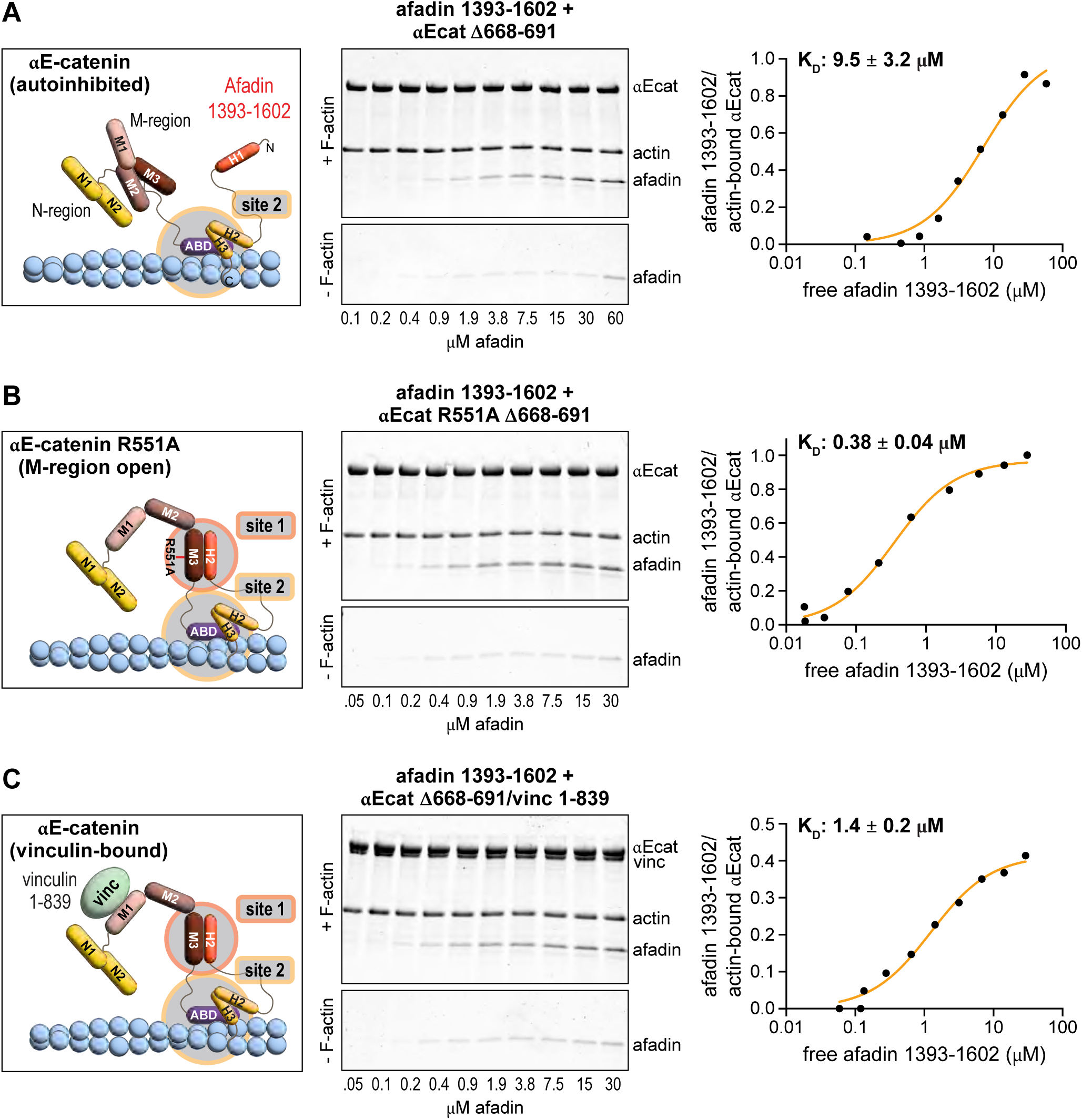
M-region opening determines afadin binding to actin-bound αE-catenin. **(A-C)** Schematic drawings and actin cosedimentation assays. Representative SDS-PAGE of pellets (+F-actin, top gel), background afadin pelleting (-F-actin, bottom gel), and binding curves of afadin binding to actin-associated αE-catenin (A), αE-catenin R551A (B), and αE-catenin in the presence of vinculin (C). Pelleting experiments were performed with β-catenin 78-151/αE-catenin Δ668-691 or β-catenin 78-151/αE-catenin R551A Δ668-691 complex. Schematic drawings indicate the accessible afadin binding sites. K_D_ values represent the average of two independent measurements.

The addition of vinculin 1-839 to autoinhibited αE-catenin Δ668-691-decorated filaments resulted in an approximately seven-fold increase in afadin affinity (K_D_ = 1.4 μM; Fig. 4, C) compared to αE-catenin Δ668-691 decorated filaments without vinculin 1-839 (Fig. 4, A), supporting our observation that vinculin binding stabilizes the open M-region conformation and unmasks the M3 domain binding site for afadin. In summary, afadin binding to the M-region is more than an order of magnitude stronger than binding to the actin-bound ABD, indicating that M-region opening is the principal determinant of afadin binding to αE-catenin.

### The afadin CC is necessary and sufficient for junctional localization in cardiomyocytes but dispensable in MDCK cells

Our biochemical experiments revealed that afadin binds to the open αE-catenin M-region with sub-micromolar affinity. Previous work from our group showed that the open αE-catenin M-region is required to recruit endogenous afadin to cardiomyocyte AJs (Merkel et al., 2019). However, αE-catenin is not required for afadin junctional localization in MDCK cells (Kuno et al., 2025), suggesting distinct mechanisms of recruitment. To determine whether afadin 1393-1602 mediates afadin localization to cardiomyocyte AJs, we expressed EGFP-tagged full-length afadin (afadin FL), 1393-1602 fragment, afadin lacking this region (afadin Δ1393-1602), or EGFP alone in cells. We then quantified junctional enrichment as the ratio of EGFP fluorescence at cell-cell contacts to cytoplasm in fixed cells (Fig. 5, A-D, E). Afadin FL was strongly enriched at cardiomyocyte junctions (EGFP contact/cytoplasm ratio = 1.90 ± 0.23), consistent with previous reports of endogenous afadin localization at cardiomyocyte cell-cell contacts (Merkel et al., 2019). Afadin 1393-1602 also showed significant junctional enrichment (ratio = 1.41 ± 0.06), but not as strongly as full-length afadin, demonstrating that the afadin CC is sufficient for junctional targeting. In contrast, afadin Δ1393-1602 showed no junctional enrichment (ratio = 0.77 ± 0.03), indistinguishable from EGFP alone (ratio = 0.70 ± 0.003), demonstrating that the 1393-1602 region is necessary for junctional targeting of afadin in cardiomyocytes. To further refine the minimal region required for localization, we tested a series of sub-fragments within 1393-1602 (Fig. 5, K). The shorter 1416-1564 fragment retained junctional localization, consistent with our *in vitro* binding results. In contrast, smaller fragments containing either helix H1 (1406-1447) or helices H2 and H3 (1520-1587, 1520-1607) were cytoplasmic, indicating that neither the M3-nor the ABD-binding helices alone were sufficient for junctional recruitment. In summary, these data show that afadin 1393-1602 is both necessary and sufficient for afadin recruitment to cardiomyocyte AJs.

**Figure 5.**
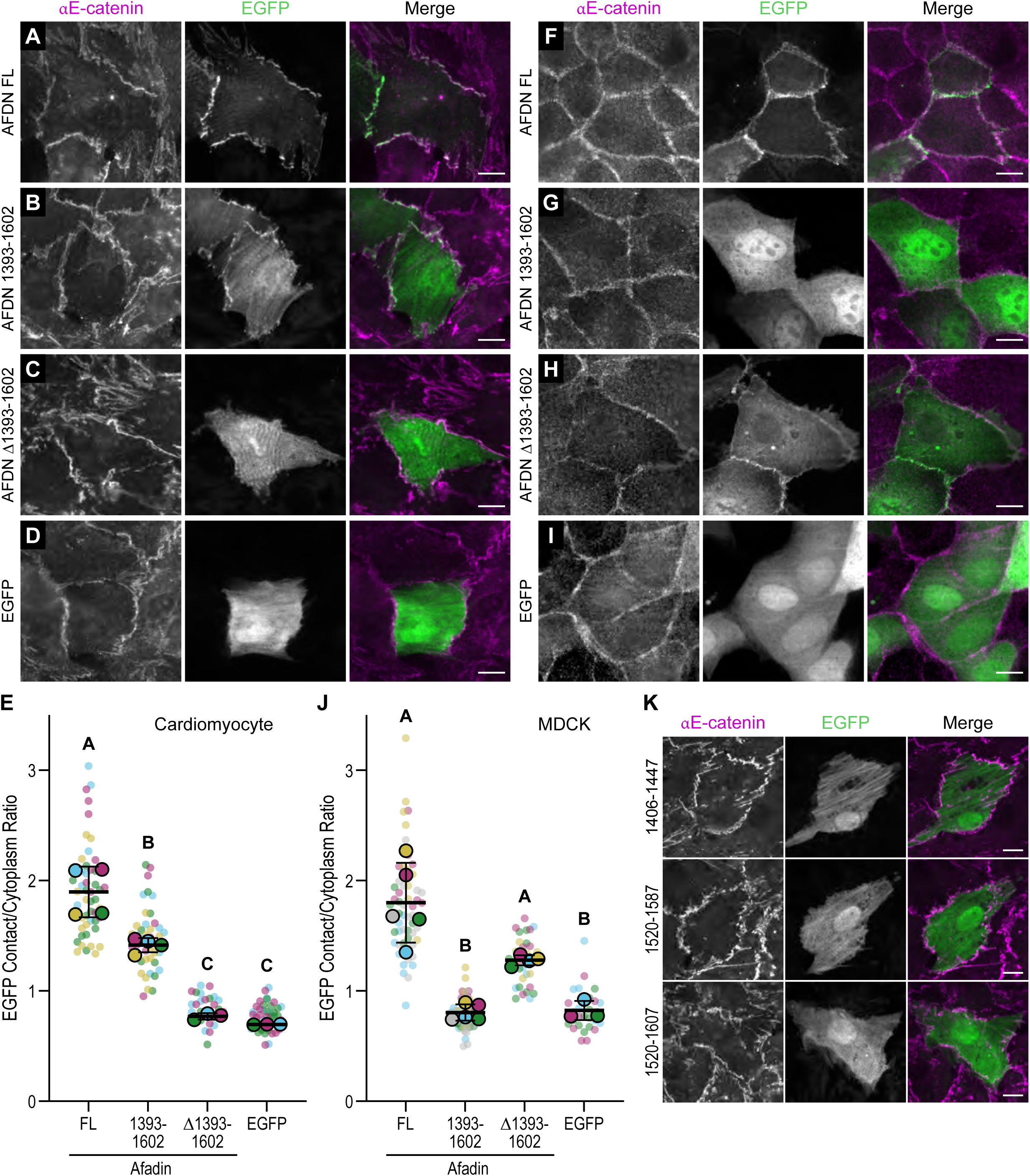
The αE-catenin-binding 1393-1602 region confers afadin junctional localization in cardiomyocytes but not MDCK cells. **(A-D)** Cultured neonatal mouse cardiomyocytes were transfected with EGFP-tagged afadin full-length (AFDN FL, A), AFDN 1393-1602 (B), AFDN Δ1393-1602 (C), or EGFP (D) alone, fixed, and stained for αE-catenin. Individual EGFP (green), αE-catenin (magenta), and merge channels are shown. Scale bars, 10 µm. **(E)** EGFP contact/cytoplasm ratio for afadin constructs expressed in cardiomyocytes. Data are presented as a SuperPlot (Lord et al., 2020). Small circles represent individual images color-coded by biological replicate; large, outlined circles indicate the replicate means. Error bars show mean ± SD. Letters indicate statistically distinct groups by one-way ANOVA with Tukey’s multiple comparisons test (p < 0.05); groups sharing the same letter are not significantly different. **(F-I)** MDCK cells were transfected with EGFP-tagged AFDN FL (F), AFDN 1393-1602 (G), AFDN Δ1393-1602 (H), or EGFP (I) alone, fixed, and stained for αE-catenin. Individual EGFP (green), αE-catenin (magenta), and merge channels are shown. Scale bars, 10 µm. **(J)** EGFP contact/cytoplasm ratio for afadin constructs expressed in MDCK cells, plotted as in E. **(K)** Cardiomyocytes transfected with EGFP-tagged afadin 1406-1447, 1520-1587, or 1520-1607 alone, fixed, and stained for αE-catenin. Individual EGFP (green), αE-catenin (magenta), and merge channels are shown. Scale bars, 10 µm.

Previously, it was shown that afadin CC localized to bicellular junctions in the mammary epithelial line Eph4 (Gong et al., 2025; Sakakibara et al., 2020), where αE-catenin is under sufficient load to relieve M-region autoinhibition (Yonemura et al., 2010). In contrast, afadin CC is not required for apical junctional localization in MDCK cells (Kuno et al., 2025). Under these conditions, vinculin is poorly recruited to MDCK cell-cell junctions (le Duc et al., 2010), consistent with them being low-load. In epithelial cells, afadin is postulated to localize to AJs primarily through interactions between its N-terminal FHA and PDZ domains and the transmembrane adhesion molecule nectin (Mandai et al., 1997; Takai and Nakanishi, 2003). To test whether afadin’s targeting requirements are conserved at low-load junctions, we expressed the same constructs in subconfluent, unpolarized MDCK monolayers, conditions expected to impose relatively low junctional tension (Fig. 5, F-I, J). As expected, afadin FL localized strongly to MDCK cell-cell junctions (ratio = 1.80 ± 0.36). Afadin Δ1393-1602 also retained junctional localization (ratio = 1.28 ± 0.05), indicating that the αE-catenin-binding region is dispensable for afadin targeting in MDCK cells. In contrast, afadin 1393-1602 failed to localize to MDCK junctions (ratio = 0.80 ± 0.07), remaining diffuse in the cytoplasm comparable to EGFP (ratio = 0.82 ± 0.09). These results revealed that in MDCK cells, αE-catenin binding was not required for afadin junctional targeting, consistent with recent work (Kuno et al., 2025). The failure of afadin 1393-1602 to target AJs suggests that the αE-catenin M-region is closed and that αE-catenin is under low load (tested in Fig. 8). We speculate that other afadin domains, such as the PDZ domain, regulate afadin localization under low-load conditions in epithelia.

### Afadin forms a dynamic biomolecular condensate at cardiomyocyte AJs

At MDCK cell–cell junctions, afadin forms biomolecular condensates that are sensitive to disruption by aliphatic alcohols such as 1,6-hexanediol (Kuno et al., 2025). To test whether afadin at cardiomyocyte junctions shows similar condensate-like properties, we treated cardiomyocytes with 5% 1,6-hexanediol or 5% 2,5-hexanediol, a structural isomer control that is markedly less effective at dissolving condensates (Kroschwald et al., 2015). 1,6-hexanediol treatment selectively and significantly reduced endogenous afadin from cardiomyocyte AJs relative to 2,5-hexanediol, whereas αE-catenin localization was unaffected (Fig. 6, A and B), consistent with afadin forming a condensate. We then tested whether EGFP-tagged afadin FL and afadin 1393-1602 were sensitive to 1,6-hexanediol. Treatment of transfected cardiomyocytes with 1,6-hexanediol, but not 2,5-hexanediol, disrupted the junctional localization of both constructs (Fig. 6, C and D). The 1,6-hexanediol sensitivity of EGFP-tagged afadin FL, which retains the complete IDR, is consistent with this construct participating in junctional condensates along with endogenous afadin. The 1393-1602 fragment contains the αE-catenin binding interface and a ∼ 70 aa IDR (the region between H1 and H2/H3), though it lacks the larger flanking IDRs (Fig. 1, B). The fragment’s localization was likewise 1,6-hexanediol-sensitive, though because 1,6-hexanediol also perturbs folded-domain interactions (Duster et al., 2021) this is consistent with, but not diagnostic of, condensate formation, as disruption of the αE-catenin-binding interface could produce the same effect.

**Figure 6.**
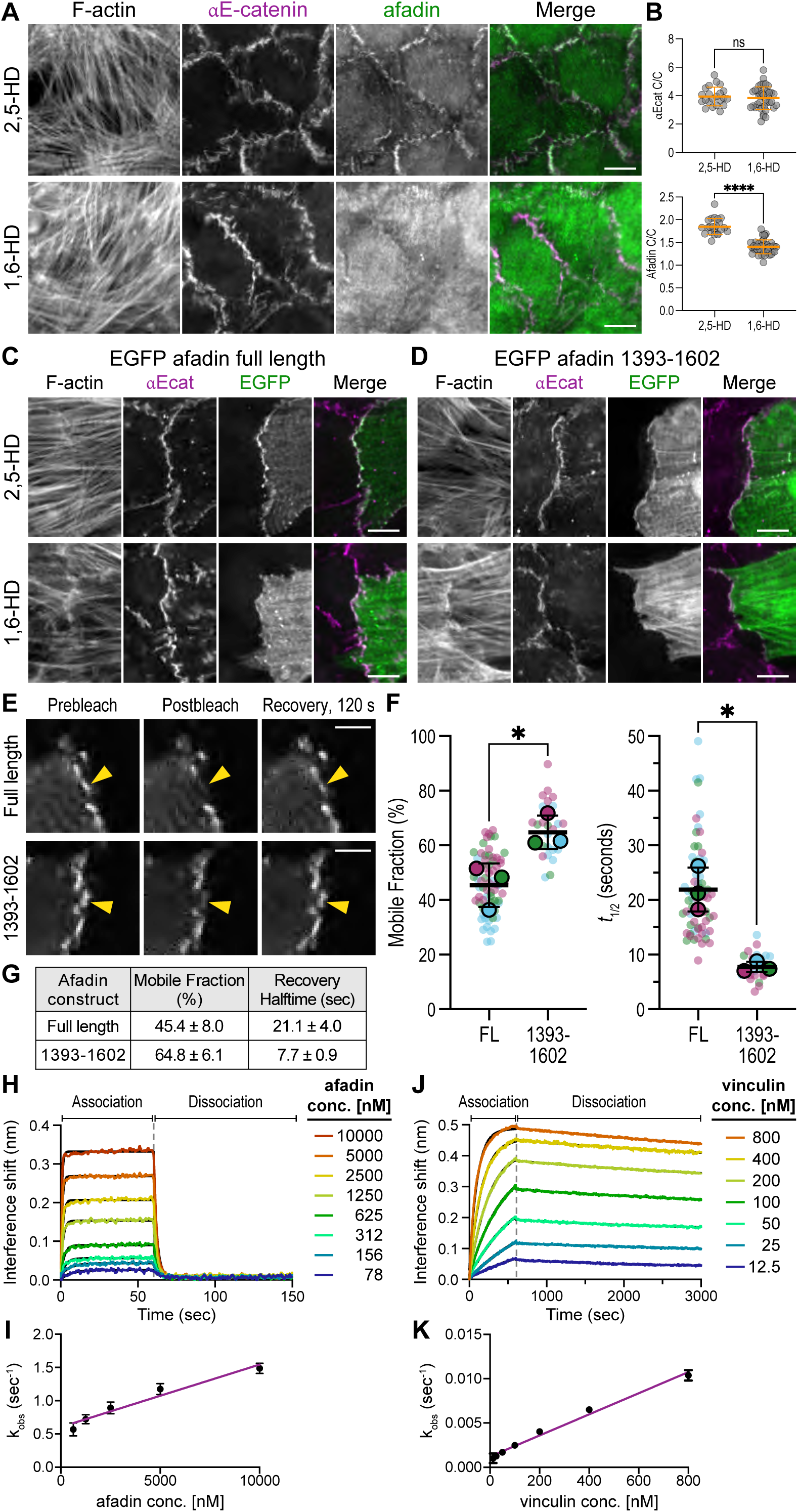
Full-length afadin exhibits condensate-like dynamics at cardiomyocyte junctions. **(A)** Representative immunofluorescence images of neonatal cardiomyocytes following treatment with 5% 2,5-hexanediol (top row) or 5% 1,6-hexanediol (bottom row) for 5 min (bottom). Cells were stained for F-actin (left), αE-catenin (second column), and afadin (third column). Merge shows αE-catenin (magenta) and afadin (green). Scale bars, 10 µm. **(B)** Quantification of afadin and αE-catenin contact/cytoplasmic (C/C) ratios in treated cells from (A). Error bars show mean ± SD. Unpaired t-test, ns = not significant, ****p < 0.0001. **(C, D)** Representative immunofluorescence images of neonatal cardiomyocytes transfected with EGFP-tagged afadin full length (C) or afadin 1393-1602 (D) and treated with 5% 2,5-hexanediol (top row) or 5% 1,6-hexanediol (bottom row) for 5 min. Cells were fixed and stained for F-actin and αE-catenin (αEcat). Merge shows αE-catenin (magenta) and EGFP (green). Scale bars, 10 µm. **(E)** Representative FRAP image sequences of neonatal cardiomyocytes expressing EGFP-tagged afadin full length (top) or afadin 1393-1602 (bottom). Prebleach, immediate postbleach, and recovery (t = 120 s) images are shown. Yellow arrowheads indicate the bleached region of interest. Scale bar, 5 µm. **(F)** Mobile fraction (% recovery) and recovery half-time (*t*_½_) of afadin full length (FL) and afadin 1393-1602, shown as a SuperPlot. Small circles indicate individual FRAP regions color-coded by biological replicate; larger, outlined circles mark the mean of each biological replicate. Horizontal bars define the mean ± SD of the biological replicate means. n = 71 FRAP regions (FL) and 26 FRAP regions (1393-1602), from N = 3 biological replicates per construct. *p < 0.05, unpaired two-tailed Welch’s t-test on biological replicate means. **(G)** Summary of mobile fraction and recovery half-time values. **(H-K)** αE-catenin binding to afadin 1393-1602 and vinculin 1-839 measured by BLI. BLI sensorgram of β-catenin 78-151/αE-catenin R551A complex binding to afadin 1393-1602 (H) or vinculin 1-839 (J) at shown concentrations. The data were fit with a single-phase association and dissociation kinetic model (black lines), yielding the observed association rate (k_obs_) and the dissociation rate constant (k_off_). The association rate constant (k_on_) was determined by fitting the concentration-dependent observed association rates (k_obs_) to a linear model (I, K).

Afadin is highly dynamic at epithelial junctions, consistent with it forming a biomolecular condensate (Kuno et al., 2025). To measure afadin dynamics at cardiomyocyte AJs, we performed fluorescence recovery after photobleaching (FRAP) on cardiomyocytes expressing EGFP-tagged afadin FL and 1393-1602 (Fig. 6, E). Individual recovery curves were fit to a single exponential model to extract mobile fraction and recovery half-time (Fig. 6, F and G). Afadin FL dynamics (mobile fraction = 45.4 ± 8.0%; *t*_½_ = 21.1 ± 4.0 s) at cardiomyocyte AJs were notably similar to afadin FL properties reported in MDCK cells (mobile fraction = ∼50%; *t*_½_ = 25.3 ± 4.9 s (Kuno et al., 2025)), and were consistent with liquid-like properties. The afadin 1393-1602 fragment exhibited a significantly larger mobile fraction and shorter recovery half-time than afadin FL (64.8 ± 6.1% *t*_½_ = 7.7 ± 0.9 s). Together with the 1,6-hexanediol results, this suggests that the afadin CC is forming a weak, highly dynamic condensate, though we cannot exclude that the isolated CC is not forming a condensate and is simply exchanging rapidly at the M-region (see below). However, the ability of afadin CC to form a weak condensate could explain why H1 alone, which lacks an IDR and ability to condense, is insufficient for junctional enrichment (Fig. 5) despite containing the primary binding site for αE-catenin M3 (Fig. 4). Consistent with this, a mutant version of the afadin CC—in which the αE-catenin binding site 2 (H2 and H3) was disrupted but H1 and the small IDR were retained—localized to junctions in Eph4 cells (Gong et al., 2025).

### Afadin and vinculin interact with αE-catenin with markedly different binding kinetics

We compared afadin dynamics to our previously measured vinculin FRAP properties at cardiomyocyte AJs (Li et al., 2025). The afadin FL mobile fraction was similar to the vinculin mobile fraction (45.4% versus 47.8%; (Li et al., 2025)); however, the recovery halftimes differed markedly: 21 seconds for afadin (Fig. 6, G) versus 136 seconds for vinculin (Li et al., 2025), a >6-fold difference. We asked whether these distinct dynamics reflect, in part, inherent differences in how these adaptor proteins associate with αE-catenin. To examine the binding kinetics directly, we used biolayer interferometry (BLI) to measure the kinetic rates of afadin 1393-1602 and vinculin binding to αE-catenin R551A. αE-catenin R551A bound afadin 1393-1602 with fast association and dissociation kinetics (Fig. 6, H). We observed a significant discrepancy between the K_D_ derived from the kinetic rates (5.4 μM) and the K_D_ determined by thermodynamics (ITC, K_D_ = 0.41 ± 0.01 μM, Fig. 2, E), which suggests that the kinetic measurements were confounded by mass transport limitations (Schuck and Zhao, 2010). This precluded the exact determination of the true kinetic rates for the αE-catenin R551A – afadin interaction (Fig. 6, I). In contrast, vinculin exhibited markedly slower binding kinetics (k_on_: 1.19 × 10⁻⁵ ± 4.7 × 10⁻⁷ nM⁻¹s⁻¹, k_off_: 0.00022 ± 0.00004 s⁻¹; Fig. 6, J and K). The K_D_ derived from the kinetic measurement (k_off_/k_on_, 18 nM) was in close agreement with the K_D_ previously determined by ITC (αE-catenin M1-M2 and vinculin 1-839: 15 nM, (Choi et al., 2012)). In conclusion, afadin and vinculin bind αE-catenin with strikingly different kinetics both *in vitro* and in cardiomyocytes: afadin forms a highly dynamic complex and vinculin a stable, long-lived complex.

### Mechanical load maintains αE-catenin-mediated afadin recruitment

Mechanical load is required to open the αE-catenin M-region (Yao et al., 2014; Yonemura et al., 2010), and our studies showed that afadin 1393-1602 binding to αE-catenin requires the M-region to be open (Fig. 2, C and D). Consistent with this, we previously reported that prolonged blebbistatin treatment reduces endogenous afadin localization at cardiomyocyte AJs (Merkel et al., 2019). To determine whether the αE-catenin-binding region of afadin specifically requires mechanical load for junctional recruitment, we treated cardiomyocytes expressing afadin FL or 1393-1602 with either blebbistatin or DMSO for 10, 20, or 30 min to inhibit myosin-dependent contractility and tension (Fig. 7, A and B). We then quantified EGFP-afadin junctional enrichment (Fig. 7, C and D). In DMSO-treated cells, both afadin FL and 1393-1602 remained stably localized at junctions across the 30-min time course. In contrast, blebbistatin treatment progressively reduced junctional localization of both constructs over time. For afadin FL, junctional enrichment was significantly reduced by 20 min (p = 0.007) and 30 min (p = 0.017) of blebbistatin relative to DMSO, though junctional localization remained well above EGFP at 30 min (ratio = 1.14 ± 0.09, above the EGFP baseline ratio 0.7 ± 0.003). The 1393-1602 fragment showed a similar yet stronger response to blebbistatin: by 20 min, junctional enrichment was significantly reduced (p = 0.019), and the contact/cytoplasm ratio had reduced to EGFP alone (ratio = 0.84 ± 0.08; p = 0.19). At 30 min, junctional enrichment remained significantly reduced relative to DMSO (p = 0.005) and statistically indistinguishable from EGFP (p = 0.51). These results demonstrate that actomyosin contractility is required to maintain afadin at cardiomyocyte AJs. Afadin 1393-1602 was more sensitive to blebbistatin than afadin FL, consistent with it forming a weaker condensate that lacks multivalent interactions necessary for condensate stability. Notably, afadin FL was partially retained at junctions even after the fragment was fully lost post blebbistatin treatment. This suggests that the IDR-mediated condensate sustains junctional afadin beyond the timescale of M-region reclosure. Together with the 1,6-hexanediol sensitivity demonstrated in Fig. 6, these results support a two-step model in which afadin is recruited to junctions through force-dependent binding to the open αE-catenin M-region and subsequently sustained by a biomolecular condensate.

**Figure 7.**
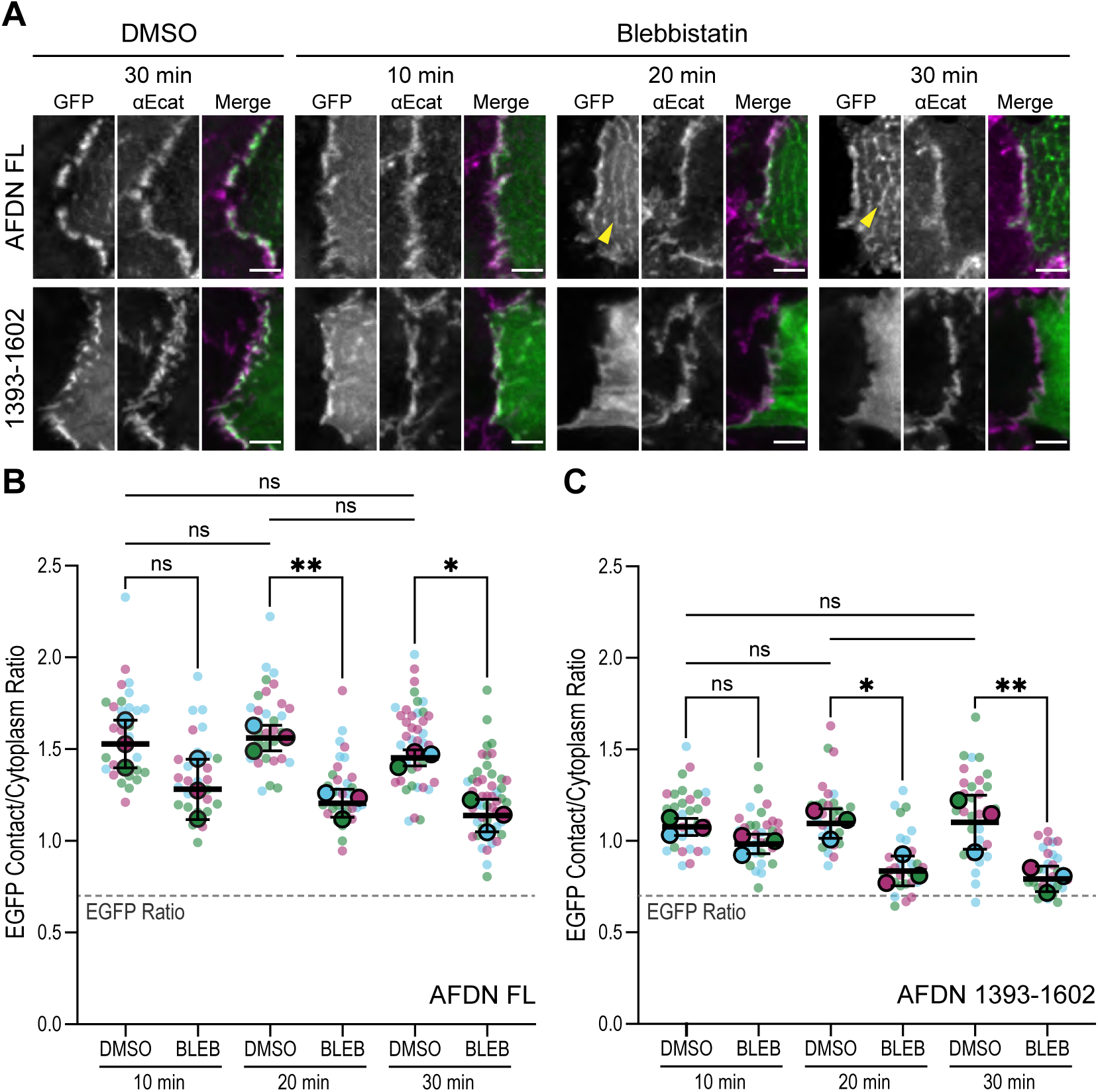
Actomyosin tension regulates afadin recruitment to cardiomyocyte AJs. **(A)** Cardiomyocytes transfected with EGFP-tagged AFDN FL (top row) or AFDN 1393-1602 (bottom row) were cultured for 48–72 h, incubated for 10, 20, or 30 min in DMSO or 50 µM blebbistatin, fixed, and stained for αE-catenin. Individual EGFP (green), αE-catenin (magenta), and merge channels are shown. Yellow arrowheads mark EGFP-AFDN FL localization to Z-discs. Scale bars, 5 µm. **(B, C)** EGFP contact/cytoplasm ratios of AFDN FL (B) and AFDN 1393-1602 (C) following DMSO or blebbistatin treatment. Small circles represent individual cells color-coded by biological replicate; larger, outlined circles indicate the replicate means. Error bars show mean ± SD. Dashed gray line indicates the mean EGFP control ratio (from Fig. 5E) for reference. All pairwise EGFP afadin comparisons were performed by one-way ANOVA with Tukey’s multiple comparisons test. ns = not significant, *p < 0.05, **p < 0.01. Comparisons to EGFP alone (Fig. 5E) were performed by one-way ANOVA with Dunnett’s multiple comparisons. All ratios in (B, C) were significantly different from EGFP except AFDN 1393-1602 BLEB at 20 min (p = 0.19) and 30 min (p = 0.51).

### αE-catenin M-region opening enables cooperative co-recruitment of afadin and αE-catenin

Our localization studies showed that afadin 1393-1602 fails to localize to MDCK junctions (Fig. 5), suggesting that endogenous αE-catenin is in a low load, autoinhibited state in MDCK cells. To determine whether αE-catenin M-region opening is sufficient to recruit afadin 1393-1602 to epithelial junctions, we co-expressed afadin constructs with αE-catenin R551A in MDCK cells. MDCK cells were co-transfected with wild-type or R551A mCherry-αE-catenin along with EGFP-tagged afadin FL, afadin 1393-1602, or EGFP alone, and the contact/cytoplasm ratios of EGFP, mCherry-αE-catenin, and endogenous β-catenin were quantified (Fig. 8, A and B). β-catenin localization was unchanged across all conditions (Fig. 8, D), consistent with results in cardiomyocytes (Fig. 8, E). Afadin FL localized to junctions in both wild-type and R551A αE-catenin backgrounds comparably (ratio = 1.52 and 1.57), demonstrating that opening of the αE-catenin M-region did not enhance afadin FL recruitment, consistent with αE-catenin-independent recruitment of FL in MDCK cells (Fig. 5; (Mandai et al., 1997) (Takai and Nakanishi, 2003)). Strikingly, the 1393-1602 fragment, which was cytoplasmic in cells expressing wild-type αE-catenin (ratio = 0.83), localized strongly to junctions when co-expressed with R551A αE-catenin (ratio = 1.93, p < 0.05; Fig. 8, D), comparable to afadin FL. Furthermore, junctional recruitment of the fragment was accompanied by a dramatic enrichment of R551A αE-catenin at junctions (ratio = 4.43, compared to 0.97 in cells expressing R551A with afadin FL; p < 0.05). This mutual dependence indicates that the open αE-catenin M-region and the afadin CC fragment co-recruit one another cooperatively. αE-catenin enrichment required both an open M-region and the isolated afadin fragment: wt αE-catenin (M-region closed) was not enriched by the fragment, and R551A αE-catenin was not enriched when co-expressed with afadin FL (Fig. 8, C and B). Thus, neither M-region opening nor afadin expression alone was sufficient—only the combination of an open M-region and an accessible afadin CC region promoted αE-catenin enrichment at junctions. That afadin FL, which contains the same αE-catenin-binding region as the fragment, did not promote αE-catenin enrichment suggests that the availability of the afadin CC differs by cellular context (see Discussion).

**Figure 8.**
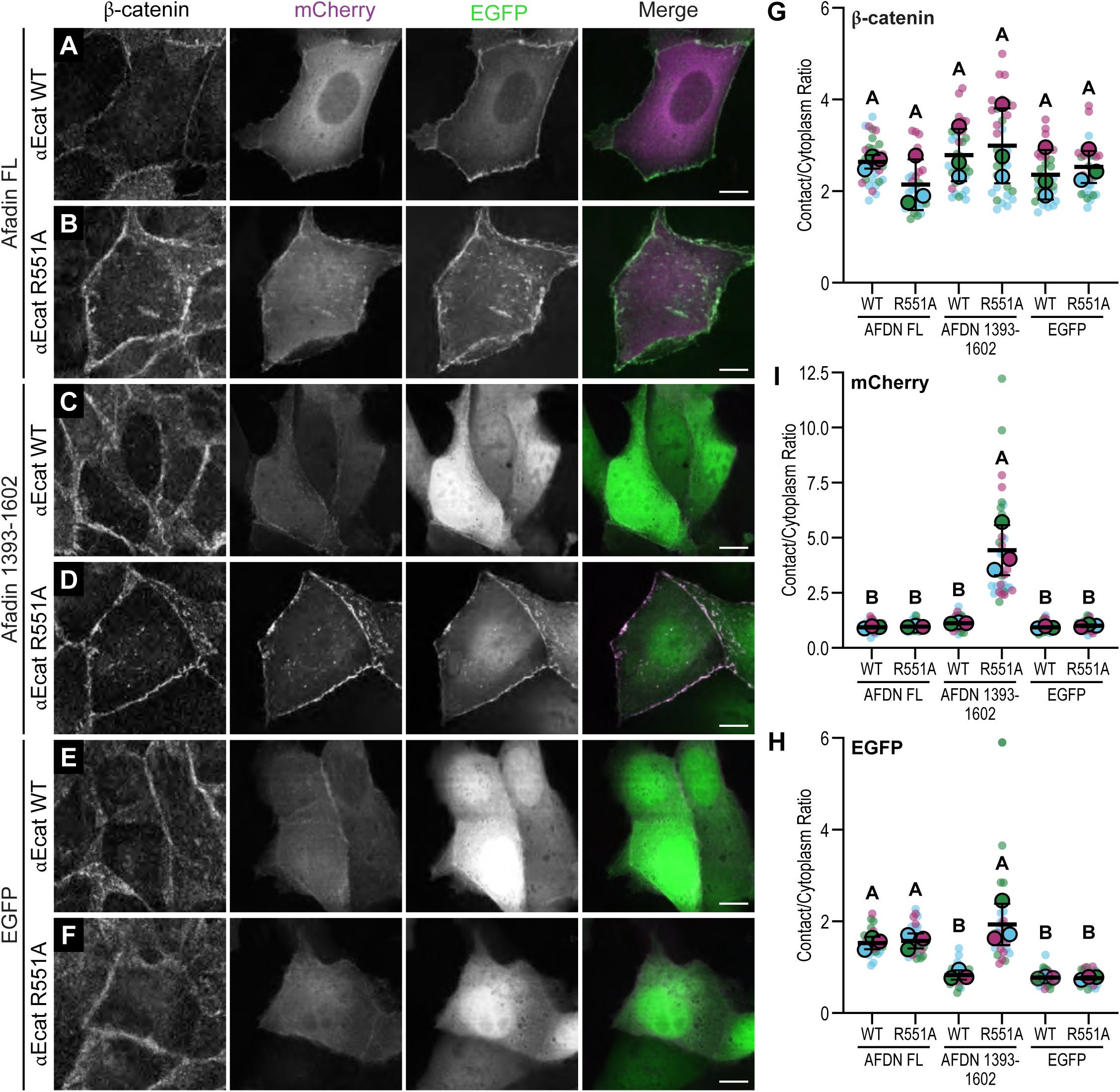
αE-catenin R551A drives cooperative recruitment of minimal but not full-length afadin in MDCK cells. **(A-F)** Representative confocal images of MDCK cells co-expressing mCherry-tagged wild-type or R551A αE-catenin together with EGFP-tagged AFDN FL (A, B), AFDN 1393-1602 (C, D), or EGFP alone (E, F). Cells were fixed and stained for endogenous β-catenin. EGFP (green) and mCherry (magenta) channels are shown individually and as a merge; β-catenin is shown separately. Scale bar, 10 µm. **(G-I)** Contact/cytoplasm ratios of endogenous β-catenin (G), and exogenous mCherry (H) and EGFP (I) in transfected MDCK cells. Small circles represent individual cells color-coded by biological replicate; larger, outlined circles indicate the replicate means. Error bars show mean ± SD. Letters indicate statistically distinct groups by one-way ANOVA with Tukey’s multiple comparisons test (p < 0.05); groups sharing the same letter are not significantly different.

## Discussion

To maintain proper adhesive function in dynamic, high-load environments, the AJ must withstand large actomyosin forces while also recognizing and responding to changes in those forces. It is well established that adaptor proteins are recruited to αE-catenin to strengthen AJs under load. Vinculin is recruited to force-activated αE-catenin, creating another F-actin binding interface that helps anchor the AJ to F-actin. Here we show that in cardiomyocytes afadin is recruited to force-activated αE-catenin through force-gated condensate formation. The afadin condensate forms in response to the mechanical state of αE-catenin, enriching afadin at the AJ and promoting cooperative αE-catenin binding and AJ reorganization.

### Afadin recruitment to cardiomyocyte AJs requires activated αE-catenin

Previous work established that the afadin CC region binds directly to the αE-catenin M-region (Maruo et al., 2018; Sakakibara et al., 2020) and, more recently, the αE-catenin ABD (Gong et al., 2025). Here we define these interactions and measure their affinities. The afadin CC region (1393–1602) contains two structurally separate interfaces: an extended helix (H1, 1406–1446) that binds the M-region, and a helix-loop-helix motif (H2 and H3, 1520–1581) that binds the ABD. The H1 interface maps primarily to M3. Recent cryo-EM work shows that H2 and H3 contact the ABD and F-actin simultaneously (Gong et al.). We show that afadin binds the open M-region with modest affinity (0.3–0.6 μM) and the actin-engaged ABD with lower affinity (∼10 μM). In cosedimentation assays with αE-catenin R551A decorated actin filaments, afadin bound with an apparent affinity (K_D_ = 0.38 μM) essentially identical to the interaction of the isolated M1-M3 R551 measured by ITC (M1–M3 R551A, K_D_ = 0.41 μM, Fig. 2). Based on these experiments, the M-region supplies effectively all of the measured affinity. Afadin binds to αE-catenin open M-region with fast kinetics (Fig. 6) and is also highly dynamic at junctions (FRAP t_½_ = 21 s for full-length protein and 8 s for the isolated CC fragment, 1393–1602). Because afadin’s interactions with αE-catenin are modest and rapidly exchanging, sustained engagement requires local afadin concentrations well above those expected for a diffusely distributed cytosolic protein. We propose that condensate formation provides this local concentration.

### A force-gated condensate concentrates and sustains afadin

Full-length afadin undergoes liquid–liquid phase separation to form condensates in MDCK cells (Kuno et al., 2025). In cardiomyocytes, junctional afadin displays properties consistent with condensate formation. First, junctional afadin localization was disrupted by 1,6-HD but not 2,5-HD, the isomeric control; junctional αE-catenin was unaffected by either treatment, indicating that 1,6-HD acted selectively on afadin rather than disrupting the AJ core. Second, junctional afadin exchanged rapidly (t_½_ = 21 s; Fig. 6), in contrast to the cadherin-catenin core (t_½_ = 220–360 s; (Li et al., 2019)) and vinculin (t_½_ = 136 s; (Li et al., 2025)) indicating a mobile, highly dynamic pool. Together, these observations are consistent with afadin forming a condensate at cardiomyocyte AJs. By concentrating afadin locally, condensate formation converts a modest, short-lived interaction into sustained engagement with force-activated αE-catenin (Fig. 9).

**Figure 9.**
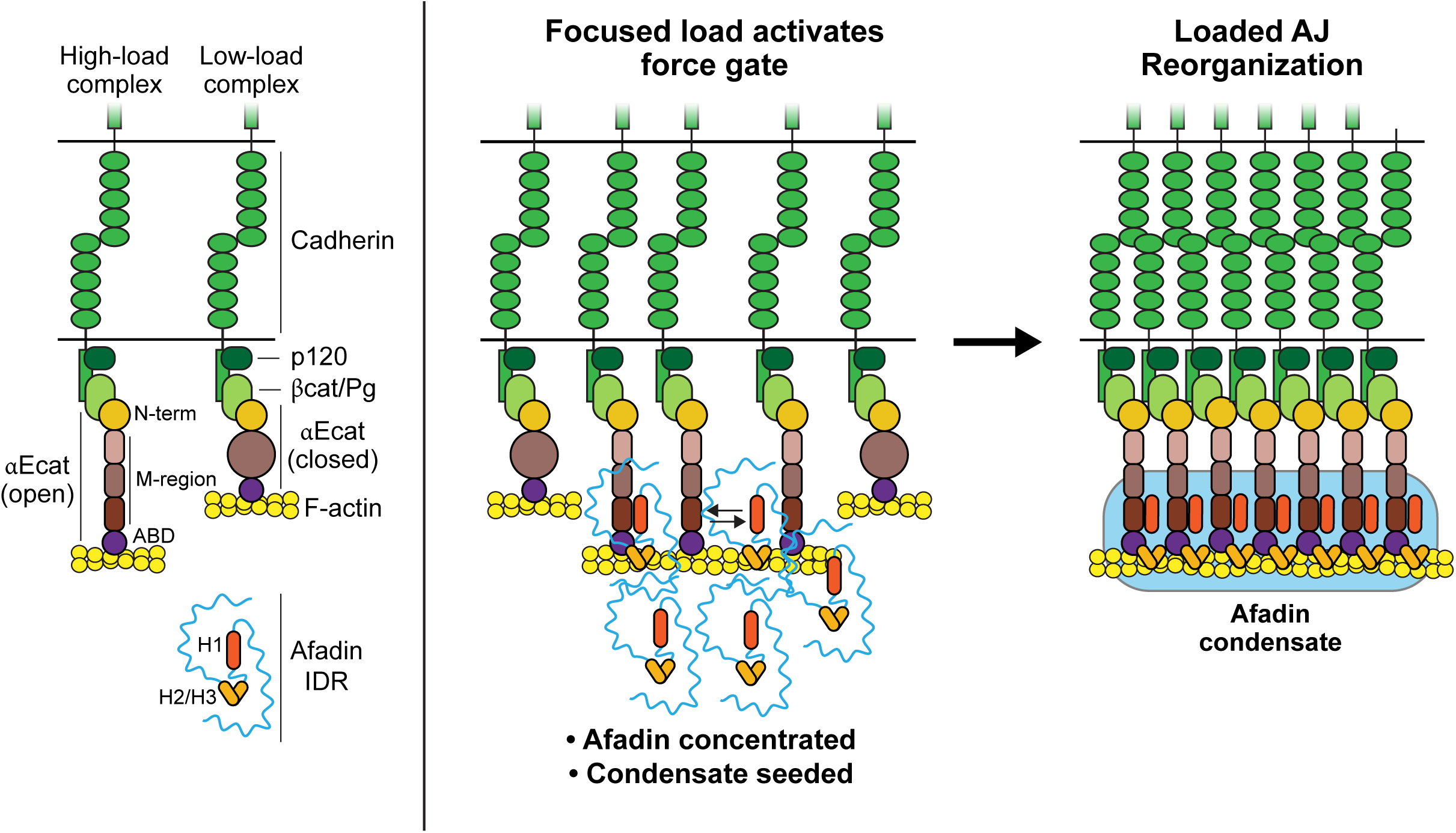
Mechanical opening of the αE-catenin M-region gates afadin condensate formation at high-load AJs in cardiomyocytes. Left: Cartoon illustration of low-load and high-load AJ complexes bound to F-actin at cell-cell contacts. Cadherin, p120-catenin, β-catenin (βcat), plakoglobin (Pg), αE-catenin (αEcat), F-actin, and afadin shown. At low-load AJs, αEcat is bound to F-actin but autoinhibited (closed). At high-load AJs, the M-region is pulled open by force. Right: Focused load at AJs activates the force gate by creating a sufficient number of high-load complexes to recruit and locally concentrate afadin to levels sufficient for condensate formation. Once formed, the afadin condensate (blue rounded rectangle) promotes AJ reorganization by clustering loaded cadherin-catenin-actin complexes, thus providing dynamic reinforcement of AJ-actin linkage.

These data help explain key features of afadin and vinculin recruitment and function in the N-cadherin–αE-catenin fusion AJ reconstitution system we described previously (Merkel et al., 2019). First, force-activated αE-catenin is required for afadin recruitment to cardiomyocyte junctions. The fusion constructs, which lacked the β-catenin binding site in N-cadherin, were expressed in N-cadherin null cardiomyocytes to rebuild AJs with defined linkages to actin. A fusion construct lacking M3 (Ncad-GFP-M1-M2) restored cell-cell junctions in N-cadherin-null cardiomyocytes and recruited vinculin in proportion to construct expression (R² = 0.40, p < 0.0001). In contrast, afadin was not recruited to this construct, with levels below those of the control (Ncad-GFP) junctions, demonstrating that αE-catenin M3 is required for afadin junctional localization in cardiomyocytes. Note that while nectins are expressed in cardiomyocytes (Satomi-Kobayashi et al., 2009), our work indicates they are insufficient to recruit afadin to cardiomyocyte cell-cell junctions on their own (Merkel et al., 2019). This contrasts with MDCK cells, where afadin is recruited to junctions independent of αE-catenin (Kuno et al., 2025). Second, in a fusion where M1 was deleted (removing the vinculin binding site) but M3 was present and exposed (Ncad-GFP-M2-ABD), afadin was recruited to junctions in the absence of vinculin. However, in contrast to vinculin, there was not a positive correlation between fusion expression and afadin recruitment (R² = 0.02, p = 0.33 across a ∼7-fold range). This is the behavior expected when recruitment is gated by a requirement beyond M3 availability: for instance, a local concentration threshold for condensate formation. Finally, selective, individual recruitment of vinculin or afadin to cardiomyocyte AJs had striking effects on junction organization. Vinculin recruitment alone was necessary and sufficient for myofibril coupling at AJs and forming junctions morphologically similar to those observed in control cells. In contrast, afadin recruitment alone was incapable of maintaining myofibril coupling and instead produced long, linear junctions (Merkel et al., 2019). Given our current results, we now posit that afadin recruitment and concomitant condensate formation promotes AJ reorganization. However, in the absence of vinculin, these junctions lack the primary link to loaded F-actin (Gong et al., 2025), and thus are unable to maintain stable connections to myofibrils.

### Junctional load determines the basis of afadin recruitment

In MDCK cells full-length afadin was not recruited to junctions by αE-catenin R551A and afadin recruitment was previously shown to not require αE-catenin (Kuno et al., 2025). In contrast, afadin CC (1393–1602) was recruited to MDCK junctions when co-expressed with αE-catenin R551A. This suggests that full-length afadin is somehow limited from engaging open αE-catenin in this epithelial context. We speculate that the route of afadin recruitment tracks junctional load. At low-load junctions, such as the bicellular contacts of MDCK monolayers, force-activated αE-catenin is scarce and afadin is recruited predominantly through nectin and other αE-catenin-independent interactions. At high-load junctions, such as cardiomyocyte AJs, force-activated αE-catenin is abundant, and recruitment is driven by its mechanical state. The reciprocal localization of the afadinΔ1393–1602 construct, which localizes to AJs in MDCK cells but is cytoplasmic in cardiomyocytes, marks these low-load and high-load routes. The CC fragment, lacking the sequences that target afadin through the nectin route, is correspondingly cytoplasmic in MDCK cells. Afadin condensates in both contexts are seeded by multivalent interactions, but the seeding interactions differ. This framework predicts that high-load junctions in epithelia, such as those in Eph4 cells, also employ the αE-catenin-gated route (Gong et al., 2025; Sakakibara et al., 2020).

### Afadin condensates promote reorganization of high-load AJs

Afadin recruitment has structural consequences for αE-catenin–actin engagement. The afadin H2/H3 helices bridge adjacent αE-catenin molecules on F-actin, promoting their cooperative clustering along the filament (Gong et al., 2025). This provides a mechanism by which force-gated binding is converted into an amplified, self-reinforcing state: once afadin is recruited to a loaded junction and locally concentrated within a condensate, it enhances cooperative binding of αE-catenin to actin filaments and facilitates the clustering of cadherin-catenin-actin linkages in high-load areas.

This mechanism also means that mechanical load across αE-catenin is transduced into the formation of a condensate that acts upon the load bearing αE-catenin–actin linkage (Fig. 9). Because this condensate layer bears no load directly, it can assemble, expand, or dissolve in response to the mechanical environment without compromising the mechanical continuity of the junction. Force-gated condensate formation therefore permits adjustment of junctional composition and organization without disrupting the load-bearing machinery itself, allowing cells to remodel adhesions while maintaining them.

## Conclusion

Changing mechanical demands in tissues require cell-cell adhesions to be both resilient and responsive. Here we describe a mechanism that meets both requirements: force-gated afadin condensate formation at mechanically activated AJs. Afadin condensates facilitate the reorganization of high-force AJs to locally concentrate loaded cadherin/catenin/actin complexes, thus providing dynamic reinforcement of AJ-actin linkage. In addition, the afadin condensate, once assembled, may outlast the mechanical input that seeded it. This decouples junctional afadin from instantaneous load, allowing the condensate to sustain a junctional state set by prior mechanical input rather than the current one. Given the prevalence of IDRs across the proteome and the growing number of adhesion adaptors shown to undergo phase separation, we speculate that force-gated condensate formation may be a general mechanism of adhesion regulation, particularly in high-load tissues such as the heart.

## Methods

### Plasmids

The rat afadin cDNA cloned into pEGFP-C1 was a kind gift from Yoshimi Takai (Nakata et al., 2007). All afadin fragments used in this study were generated by PCR using pEGFP-C1-afadin (rat) as the template. For mammalian expression studies, afadin fragments were cloned into pEGFP-C1 or pEGFP-N1 by Gibson assembly (New England Biolabs HiFi Kit). For protein expression, afadin 1416-1564 was cloned into pMAL-c6T (New England Biolabs) by Gibson assembly. All other afadin (afadin 1393-1602, afadin 1402-1451) murine αE-catenin fragments (αE-catenin R551A, αE-catenin 271-633 (M1-M3), αE-catenin 271-633 R551A (M1-M3 R551A), αE-catenin 397-906 (M2-ABD), αE-catenin 508-906 (M3-ABD), αE-catenin 508-633 (M3)), murine β-catenin 78-151 and 78-671 fragments and chicken vinculin 1-839 were cloned by restriction cloning or Gibson assembly into a modified pGEX-KG vector into which a TEV site was inserted (pGEX-TEV). The N-terminal fusion protein of afadin 1402-1451 and αE-catenin 507-633 was designed with a six amino acid linker (GGSGGS) and generated by Gibson assembly. The R551A point mutation and the Δ668-691 deletion in αE-catenin were introduced by site-directed mutagenesis (NEB Q5 site-directed mutagenesis kit). For coexpression with β-catenin, αE-catenin was cloned into a pET29a vector (Epoch Life Science).

### Protein expression and purification

Afadin 1416-1564 was expressed as tobacco etch virus (TEV) protease cleavable MBP fusion protein. All other afadin (afadin 1393-1602, afadin 1402-1451, afadin 1402-1451-αE-catenin-M3 fusion) and αE-catenin (αE-catenin R551A, αE-catenin M1-M3, αE-catenin M1-M3 R551A, αE-catenin M2-ABD, αE-catenin M3-ABD, αE-catenin M3) constructs, and vinculin 1-839 were expressed as TEV cleavable GST fusion proteins. Complexes with β-catenin (β-catenin 78-671 or β-catenin 78-151) and α-catenin were prepared by coexpressing TEV-cleavable GST-β-catenin and untagged αE-catenin (pET29a vector).

Constructs were transformed into *Escherichia coli* BL21(DE3) cells. Cultures were grown at 37°C to an OD_600_ of 0.6-0.8 and induced overnight at 18 °C with 0.5 mM isopropyl 1-thio-β-D-galactopyranoside. Cells were harvested by centrifugation, and the cell pellet was resuspended in 20 mM Tris pH 8.0, 150 mM NaCl, 1 mM DTT. After addition of protease inhibitors (0.15 μM aprotinin, 1 μM E-64 and 1 μM leupeptin final concentration) and DNase (Sigma) cells were lysed in an Emulsiflex (Avestin). The lysate was cleared by centrifugation at 40 000 x g for 30 min at 4°C and the supernatant was incubated with glutathione agarose, or in case of afadin 1416-1564 with amylose resin for 1h at 4°C. After washing with PBS containing 500 mM NaCl, 1mM DTT and 0.005% Tween 20, beads were equilibrated with TEV cleavage buffer (20mM Tris pH 8.0, 150 mM NaCl, 1mM DTT, 1mM EDTA, 10% glycerol) and incubated overnight at 4°C with TEV protease. The cleaved protein was further purified by anion exchange chromatography (Mono Q 10/100, Cytiva), or, for afadin 1416-1564, by cation exchange chromatography (Mono S 10/100, Cytiva) followed by gel filtration chromatography (Superdex 200, Cytiva) in 20 mM HEPES pH 8.0, 150 mM NaCl, 1mM DTT.

### Isothermal Titration Calorimetry

ITC titrations were performed in a MicroCal VP-ITC calorimeter (Malvern Panalytical). Purified protein in 20 mM HEPES pH 8.0, 150 mM NaCl, 1mM DTT was used at concentrations between 9-14 μM in the cell and 98-195 μM in the syringe. For titrations with αE-catenin M1-M3, αE-catenin M1-M3 R551A and αE-catenin M3-afadin fusion protein, afadin was placed in the cell and for the titration with αE-catenin M1-M3/afadin1393-1602 complex and vinculin 1-839, vinculin was in the syringe. All other titrations were performed with afadin in the syringe. The αE-catenin M1-M3 R551A/vinculin 1-839 complex was prepared by incubating the two proteins at a 1:1 molar ratio for 30 min at room temperature and subsequent purification by gel filtration chromatography. For ITC experiments with αE-catenin M1-M3 and vinculin or αE-catenin M1-M3 and afadin 1393-1602 in the cell, 10 μM αE-catenin M1-M3 and 20 μM vinculin 1-839 or 15 μM afadin was used. Titrations were performed at 25°C with 10 μl injections spaced at 240 sec. Data were analyzed with the Microcal Origin software (version 7.0). For baseline correction, the average heat change at saturation was subtracted from all data points. The fit was performed with a single-site binding model.

### AlphaFold predictions

Sequences of full-length rat afadin, afadin 1400-1458, murine αE-catenin 507-635, and the sequence of the fusion protein afadin 1402-1451-GGSGGS-αE-catenin 507-633 were used as input into AlphaFold3 (Abramson et al., 2024). Models are displayed in PyMOL (The PyMOL Molecular Graphics System, Version 3.1.5.1 Schrödinger, LLC).

### Actin cosedimentation assay

G-actin prepared from rabbit muscle (Spudich and Watt, 1971) was stored in 100 μl aliquots at -80°C. Frozen aliquots of 40 μM G-actin were thawed on ice and polymerized by adding 10x F-buffer (100 mM Tris pH 7.5, 500 mM KCl, 20 mM MgCl_2_, 10 mM ATP) and incubating at room temperature for 1h. All experiments were conducted with a single batch of F-actin for which actin polymerization efficiency was confirmed by pelleting the polymerized actin for 20 min at 140,000 x g at 4°C and subsequent analysis of supernatant and pellet by SDS-PAGE.

For sedimentation assays with β-catenin 78-151/αE-catenin Δ668-691, F-actin was diluted to 4 μM with buffer A (20 mM HEPES pH 8.0, 150 mM NaCl, 2mM MgCl_2_, 1mM DTT, 1mM EGTA, 0.5 mM ATP). A dilution series of β-catenin/α-catenin complex was prepared with 20 mM HEPES pH 8.0, 150 mM NaCl, 1mM DTT, and equal volumes of either F-actin or buffer A (control) were added to each concentration point. To determine binding of afadin 1393-1602 to actin-bound α-catenin, afadin 1393-1602 was serially diluted. An equal volume of either buffer A (control) or a solution containing 4μM actin, 10 μM β-catenin 78-151/αE-catenin Δ668-691 complex, and if included, 20 μM vinculin 1-839, was added to each concentration point. Samples were incubated for 30 min at room temperature. After centrifugation at 140,000 x g for 20 min at 4°C in a Beckman TLA 100 rotor, the supernatant was carefully removed, and the pellet was resuspended in reducing Laemmli buffer. Samples were analyzed by SDS-PAGE, and the Coomassie-stained bands were quantified on a LI-COR Odyssey (LI-COR Biosciences) scanner. A concentration series of α-catenin or afadin was used as a standard to extrapolate concentrations from band intensities. Background pelleting without actin was subtracted, and band intensities were normalized to the actin band intensity for each concentration point. The data were analyzed using GraphPad Prism 10 (GraphPad Software, La Jolla, CA). Binding curves were fitted with a one–site, specific binding model.

### Size exclusion chromatography

αE-catenin M1-M3 R551A, αE-catenin M1-M3, vinculin 1-839, and afadin 1393-1602 were diluted to 20 μM in 20 mM HEPES pH 8.0, 150 mM NaCl, 1mM DTT. Ternary complexes were formed by mixing 20 μM α-catenin (αE-catenin M1-M3 R551A or αE-catenin M1-M3), vinculin 1-839 and afadin 1393-1602 and incubation for 15 min at room temperature. Samples (100 μl) were loaded onto an analytical Superdex 200 column (Superdex 200 Increase 10/30 GL, Cytiva). Fractions were separated by SDS-PAGE and stained with Coomassie.

### Biolayer Interferometry

Biolayer Interferometry assays were performed using an Octet RED384 instrument (Sartorius). Biosensors were hydrated in BLI buffer (20 mM HEPES pH 8.0, 150 mM NaCl, 1mM DTT, 0.1 % BSA, 0.08% Tween 20) for a minimum of 20 min before use. Experiments were conducted at 25°C and 1000 rpm. Data for afadin and vinculin binding were collected at 10 Hz and 5 Hz, respectively.

β-catenin 78-151/αE-catenin R551A complex was biotinylated using EZ- link NHS- PEG4- Biotin (Thermo Fisher) at a 1:1 molar ratio. The reaction was performed in 20 mM HEPES pH 8.0, 150 mM NaCl, 1mM DTT. After incubation for 30 min at room temperature, the reaction was stopped by adding 1 M Tris pH 8.0, to a final concentration of 10 mM. Unreacted NHS-PEG4-biotin was removed using a Zeba desalting column (Thermo Fisher).

Biotinylated β-catenin 78-151/αE-catenin R551A complex diluted in BLI buffer was captured on a streptavidin-coated biosensor tip until an interference shift of 2.0 nm (afadin 1393-1602 binding assay) or 0.7 nm (vinculin 1-839 binding assay) was reached. After loading, sensors were equilibrated in BLI buffer for 120 sec and then dipped into BLI buffer to establish a baseline. Sensors were dipped into ligand diluted in BLI buffer to record association and returned to the well containing buffer to measure dissociation. Afadin association and dissociation were monitored for 60 and 90 sec, respectively. Vinculin kinetics were recorded using a 10 min association and 40 min dissociation phase. In experiments with afadin 1393-1602, repeated association/dissociation cycles were recorded using the same tip dipped into increasing ligand concentrations. For vinculin 1-839, a single tip was used for each concentration point. Empty sensor tips run in parallel were used to monitor background binding. No background binding was observed for afadin 1393-1602 or vinculin 1-839 at the used concentrations. For each assay, an association/dissociation cycle in buffer was used to correct for baseline drift.

Data preprocessing (subtraction of the association/dissociation cycle at zero ligand concentration, Y axis alignment, Inter-step correction, Savitzky-Golay filtering) was performed using the Octet data processing software 10.0 HT. Curve fitting was performed in the Prism 10 software (GraphPad) using the ‘one phase association’ and ‘dissociation-one phase exponential decay’ model to obtain k_obs_ and k_off_. k_on_ was obtained from linear regression fits of K_obs_ vs concentration plots. k_off_ was determined by averaging k_off_ obtained at different concentration points using fits with R^2^ > 0.95. Data from six (afadin) and two (vinculin) independent measurements were used to determine kinetic rate constants.

### Antibodies

Primary antibodies used for immunostaining were: anti-αE-catenin (1:100, Enzo Life Science, ALX-804-101-C100), anti-β-catenin (1:100, Cell Signaling, D10A8), anti-afadin (1:500, Sigma, A0349), anti-vinculin (1:800, Sigma V9131). Secondary antibodies used were goat anti-mouse or anti-rabbit IgG labeled with Alexa Fluor 568 or 647 (1:250, Thermo Fisher Scientific). F-actin was stained with phalloidin conjugated to Alexa Fluor 568 or 647 (1:100, Thermo Fisher Scientific).

### Cardiomyocyte isolation and culture

All animal work was approved by the University of Pittsburgh Division of Laboratory Animal Resources (IACUC protocol 25107506). Primary cardiomyocytes were isolated from P1 Swiss Webster mice as described (Ehler et al., 2013). For immunostaining, cardiomyocytes were plated onto 35 mm MatTek dishes with 10 mm insets coated with Collagen Type I (Millipore Sigma). Cardiomyocytes were plated in plating medium: 65% high glucose DMEM (Thermo Fisher Scientific), 19% M-199 (Thermo Fisher Scientific), 10% horse serum (Thermo Fisher Scientific), 5% FBS (Atlanta Biologicals), and penicillin-streptomycin (Thermo Fisher Scientific). Medium was replaced 16 hours after plating with maintenance medium: 78% high-glucose DMEM, 17% M-199, 4% horse serum, penicillin-streptomycin, 1 µM AraC (Sigma), and 1 µM Isoproterenol. Cells were transfected 24 h post-plating using Lipofectamine 2000 (Thermo Fisher Scientific). Cardiomyocytes were cultured in maintenance medium for 1-3 days prior to live cell imaging or fixation.

### MDCK cell culture

MDCK G type II cells were maintained in DMEM with 1 g/L glucose, 10% fetal bovine serum (Atlanta Biologicals), and penicillin/streptomycin. For immunostaining, MDCK cells were plated onto 22 mm coverslips coated with Collagen Type I (Millipore Sigma). Cells were transfected using Lipofectamine 2000 (Thermo Fisher Scientific). MDCK cells were fixed 48 h post-transfection.

### Immunostaining and confocal microscopy

Cells were fixed in 4% EM grade paraformaldehyde in PHEM buffer (60 mM PIPES pH 7.0, 25 mM HEPES pH 7.0, 10 mM EGTA, pH 8.0, 2 mM MgCl_2_, and 0.12 M Sucrose) or PBS-sucrose buffer (PBS, Ca^2+^, Mg^2+^ plus 0.12 M sucrose) for 10 minutes, washed twice with PBS, and then stored at 4 °C until staining. Cells were permeabilized with 0.2% Triton X-100 in PBS for 5 minutes and washed twice with PBS. Cells were then blocked for 1 hour at room temperature in PBS + 10% BSA (Sigma), washed 2X in PBS, incubated with primary antibodies in PBS + 1% BSA for 1 hour at room temperature, washed 2X in PBS, incubated with secondary antibodies in PBS + 1% for 1 hour at room temperature, washed 2X in PBS and then mounted in ProLong Glass (Thermo Fisher Scientific). All samples were cured for at least 24 hours before imaging.

For blebbistatin experiments, cardiomyocytes were treated with 50 or 100 μM blebbistatin in DMSO (both concentrations showed equivalent results) or DMSO alone in maintenance media for 10, 20, or 30 minutes. Cells were incubated at 37 °C during treatment. After incubation, cells were fixed in 4% EM grade paraformaldehyde in PBS-sucrose buffer, and labeled as described.

For the hexanediol experiments, cardiomyocytes were treated with 5% 1,6-hexanediol or 5% 2,5-hexanediol (control isomer) in maintenance media for 5 minutes. Cells were incubated at 37 °C during treatment. After incubation, cells were fixed in 4% EM grade paraformaldehyde in PBS-sucrose buffer + 0.3% Triton X-100, and labeled as described.

All cells were imaged with a 100x objective (NA 1.45) on a Nikon Eclipse Ti inverted microscope equipped with a Prairie swept-field confocal scanner, an Agilent monolithic laser launch, and an Andor iXon3 camera. The microscope was run by NIS-Elements (Nikon) imaging software. 3–5 µm Z-stacks with a step size of 200 nm were collected for all fixed samples.

### Image analysis

To measure junctional enrichment of GFP-tagged afadin proteins with catenin proteins in MDCK and cardiomyocytes, a maximum intensity projection of a 1 μm z-slab centered on cell-cell contacts was generated for analysis in ImageJ (NIH). Thresholding was used to create a mask of the αE-catenin or β-catenin channel to define the region of analysis (cell–cell contacts). The average GFP and vinculin signal intensities were measured in the masked region (contacts). Three ROIs were manually drawn within transfected cells to calculate the average cytoplasmic fluorescence signal. For each image, the average contact signal was divided by the average cytoplasmic signal to calculate the contact/cytoplasmic ratio.

### FRAP

FRAP experiments were conducted on a Nikon swept-field confocal microscope (described above) outfitted with a Tokai Hit cell incubator and a Bruker miniscanner. Actively contracting cells were maintained at 37 °C in a humidified atmosphere containing 5% CO_2_. User-defined regions along cell-cell contacts—typically 2 μm wide (parallel to the junction membrane) by 4 μm long (perpendicular to the membrane)—were bleached with a 405-nm laser, and recovery images were acquired every 2 seconds for 2 minutes. FRAP data were quantified in ImageJ (NIH). FRAP recovery curves from individual regions were background-subtracted and normalized to pre-bleach and immediate post-bleach intensities. Normalized recoveries were fit to a one-phase association model [Y = Plateau × (1 − e^(−K·X))] in GraphPad Prism, version 11, with Y₀ constrained to 0 and Plateau constrained between 0 and 1. Individual fits were excluded from downstream analysis if any of the following criteria were met: (1) the plateau parameter converged to the upper constraint boundary (≥0.99); (2) the coefficient of determination (R²) was below 0.6; (3) the fitted plateau exceeded 125% of the mean of the last four acquired timepoints, indicating extrapolation beyond the observed recovery; or (4) the fitted time constant (τ) equaled or exceeded the acquisition window (120 s), indicating that the recovery had not reached plateau within the acquired data. These exclusion criteria were pre-specified and applied identically to both constructs. Mobile fraction (plateau) and half-time of recovery [t½ = ln(2)/K] were extracted from each retained fit. For statistical comparison between constructs, per-region values were averaged within each biological replicate, and biological replicate means were compared using Welch’s t-test.

## Acknowledgements

This work is dedicated to the memory of Bill Weis (1959–2023). Bill’s structural and biochemical dissection of the cadherin–catenin complex laid much of the foundation for this study, and his influence on structural biology, biochemistry, and cell biology endures. We hope it honors his legacy. We thank Cara Gottardi and Brent Hoffman for providing critical feedback on the manuscript.

---

**Supplementary Figure 1.**
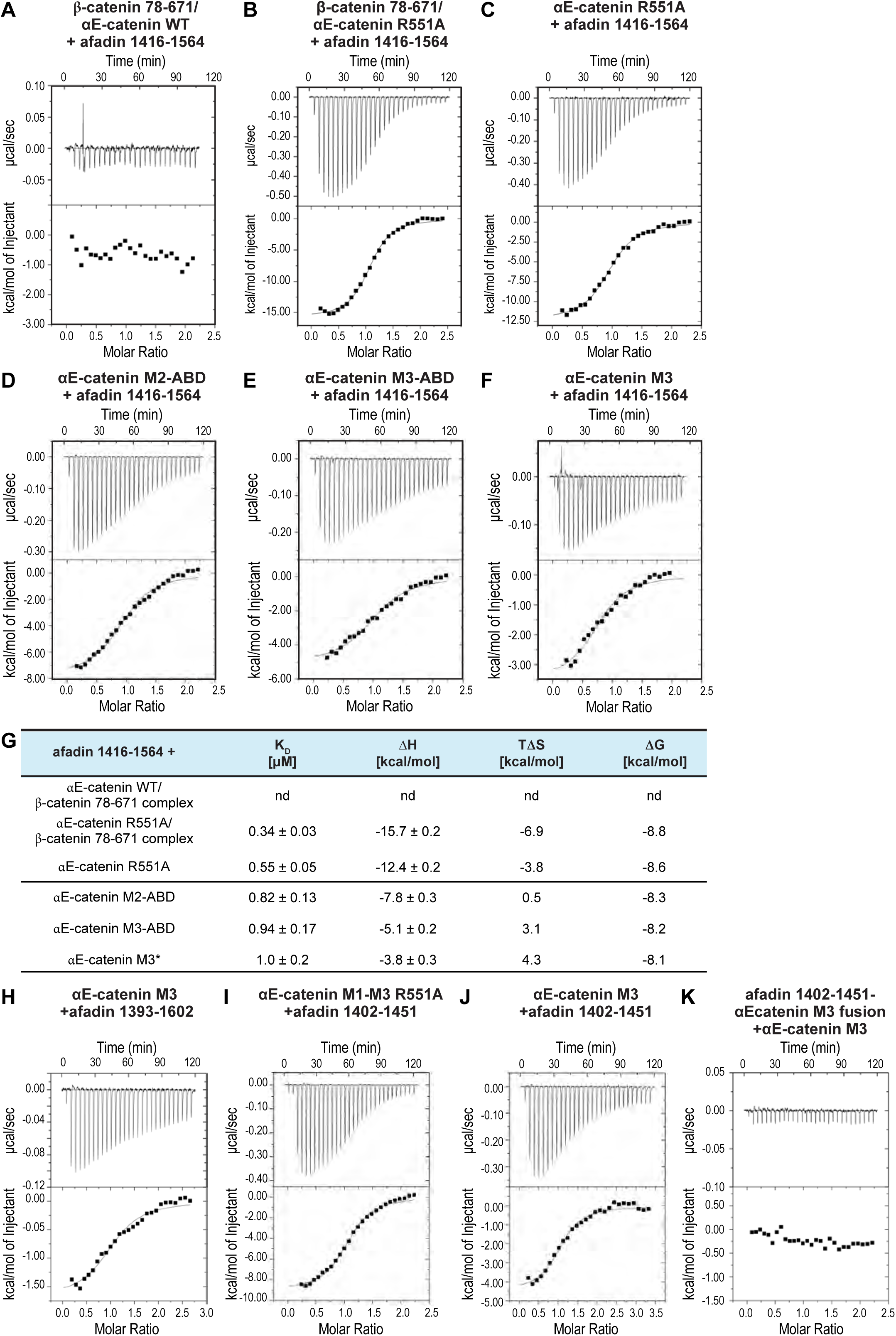
Afadin 1416-1564 and afadin 1393-1602 bind αE-catenin with comparable affinity; mapping of the afadin-αE-catenin binding site. **(A-C)** ITC traces for afadin 1416-1564 binding to wild-type (A) or R551A αE-catenin full-length in complex with β-catenin 78-671 (B) or R551A αE-catenin full-length (C). **(D-F)** ITC traces for afadin 1416-1564 binding to αE-catenin fragment M2-ABD (D), M3-ABD (E), and the isolated M3 domain (F). **(G)** Thermodynamic parameters were obtained from single measurements and derived from the fits in (A-F) with errors representing the residuals of the fit; nd = not detected. **(H-K)** Representative ITC traces for afadin 1393-1602 binding to αE-catenin M3 (H), afadin 1402-1451 binding to αE-catenin M1-M3 R551A (I), afadin 1402-1451 binding to αE-catenin M3 (J), and αE-catenin M3 binding to the afadin 1402-1451-αE-catenin M3 fusion protein (K).

**Supplementary Figure 2.**
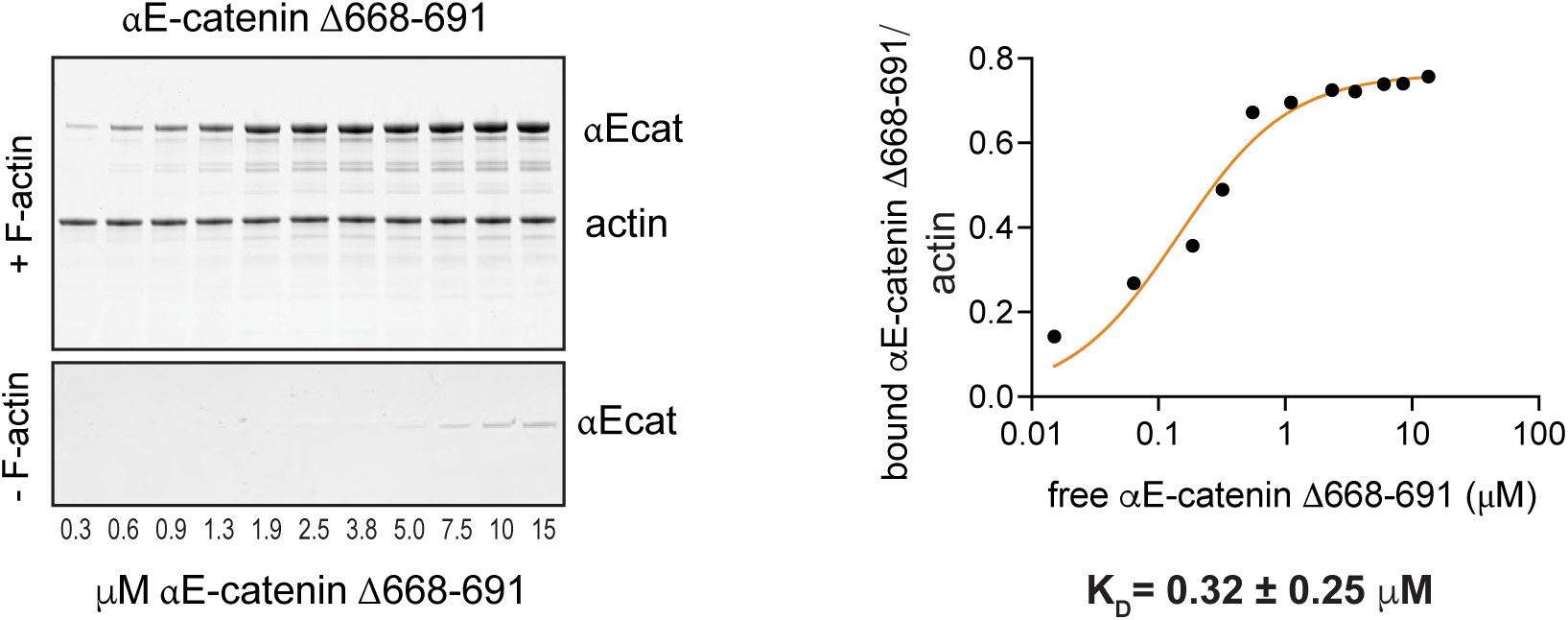
αE-catenin Δ668-691 binding to actin filaments. Actin cosedimentation assay of β-catenin 78-151/αE-catenin Δ668-691complex with representative SDS-PAGE of pellets (+ F-actin, top gel), background afadin pelleting (- F-actin, bottom gel), and binding curve. The K_D_ represents the average of two measurements.

